# DeepRank-GNN: A Graph Neural Network Framework to Learn Patterns in Protein-Protein Interfaces

**DOI:** 10.1101/2021.12.08.471762

**Authors:** M. Réau, N. Renaud, L. C. Xue, A. M. J. J. Bonvin

**Affiliations:** Computational Structural Biology Group, Department of Chemistry, Bijvoet Centre, Faculty of Science, Utrecht University, Utrecht, 3584CH, The Netherlands; Netherlands eScience Center, Science Park 140, 1098 XG, Amsterdam, The Netherlands; Center for Molecular and Biomolecular Informatics, Radboudumc, Greet Grooteplein 26-28, 6525 GA Nijmegen, The Netherlands

**Author notes:** These authors contributed equally (M. Réau, N. Renaud).

## Abstract

Gaining structural insights into the protein-protein interactome is essential to understand biological phenomena and extract knowledge for rational drug design or protein engineering. We have previously developed DeepRank, a deep-learning framework to facilitate pattern learning from protein-protein interfaces using Convolutional Neural Network (CNN) approaches. However, CNN is not rotation invariant and data augmentation is required to desensitize the network to the input data orientation which dramatically impairs the computation performance. Representing protein-protein complexes as atomic- or residue-scale rotation invariant graphs instead enables using graph neural networks (GNN) approaches, bypassing those limitations.

We have developed DeepRank-GNN, a framework that converts protein-protein interfaces from PDB 3D coordinates files into graphs that are further provided to a pre-defined or user-defined GNN architecture to learn problem-specific interaction patterns. DeepRank-GNN is designed to be highly modularizable, easily customized, and is wrapped into a user-friendly python3 package. Here, we showcase DeepRank-GNN’s performance for scoring docking models using a dedicated graph interaction neural network (GINet). We show that this graph-based model performs better than DeepRank, DOVE and HADDOCK scores and competes with iScore on the CAPRI score set. We show a significant gain in speed and storage requirement using DeepRank-GNN as compared to DeepRank.

DeepRank-GNN is freely available from https://github.com/DeepRank/DeepRank-GNN.

## 1. Introduction

Protein-protein interactions (PPIs) are essential in all cellular processes of living organisms including cell growth, structure, communication, protection and death. Adding the structural dimension to PPI is fundamental to understand normal and altered physiological processes and to propose solutions to restore them. In the past decades, a large number of isolated protein and PPI structures have been solved by experimental approaches (e.g., X-ray crystallography, nuclear magnetic resonance, cryogenic electron microscopy). The diversity and quantity of structural data recently enabled treating PPI data with machine learning approaches that were previously devoted to small molecule toxicity (Mayr *et al*., 2016), affinity (Son and Kim, 2021; Karlov *et al*., 2020; Jiménez *et al*., 2018; Ragoza *et al*., 2017) and binding mode (Torng and Altman, 2019; Morrone *et al*., 2020; Francoeur *et al*., 2020) prediction. Given the remarkable success of Convolutional Neural Network (CNN) in retrieving patterns in images (Krizhevsky *et al*., 2017), CNN approaches have been developed to learn interaction patterns in PPI interfaces (Renaud *et al*., 2021; Wang *et al*., 2020) or to assess the quality of protein structures (Baldassarre *et al*., 2021; Pagès *et al*., 2019). The uniqueness of each approach originates from the designed network architecture and importantly, the data representation and resolution. An example is MASIF (Gainza *et al*., 2020) that makes use of a high-level representation of proteins, focusing on their surface described as an ensemble of overlapping patches. The patches are fed into different CNNs in order to build relevant fingerprints that can be further used for ultra-fast interaction prediction tasks based on the complementarity or the similarity of the fingerprints. DOVE (Wang *et al*., 2020) evaluates protein-protein docking models using a 3D-CNN approach on a higher resolution -- atomic-level – representation of the interface mapped into a 3D grid. Although no exhaustive benchmark exists with state-of-the-art approaches, they both display high performance on the benchmark set used for their evaluation and hold the promise to improve over time with the availability of new data and the improvement of data storage and computation power. As the recent major advances made by Alphafold2 in predicting protein structures (Jumper *et al*., 2021) and protein multimeric states (Evans *et al*., 2021) is likely to lead to an exponential generation of multimers over years, including true and false partners, the availability of reliable quality assessment tools should become a strong ally to reach the ambitious objective of modelling of the entire interactome (Burke *et al*., 2021).

We have recently developed DeepRank (https://github.com/DeepRank/deeprank), an open-source configurable deep learning framework wrapped into a user-friendly python3 package (Renaud *et al*., 2021; Renaud *et al*., 2020). DeepRank maps atomic and residue-level features from PPIs to 3D grids and applies a customizable 3D CNN pipeline to learn problem-specific interaction patterns. DeepRank was applied to two problems where it competed with- or outperformed state-of-the-art methods, including a machine learning-based model, iScore (Geng *et al*., 2020) that actually also makes use of a graph representation and the classical energy-based scoring function implemented in HADDOCK (van Zundert *et al*., 2016).

CNNs however come with limitations: First, they are sensitive to the input PPI orientation which may require data augmentation (i.e. multiple rotations of the input data) for the network to provide consistent predictions regardless of the orientation of the PPI; second, the size of the 3D grid is pre-defined for all input data in DeepRank 0.2.0, which does not reflect the variety in interface sizes observed in experimental structures and may be problematic for large interfaces that do not fit inside the predefined grid size.

A solution to this problem is to use graph representation of PPI. A graph is defined as an ensemble of nodes (e.g.: atoms, residues) and edges (e.g.: covalent bond, contacts), and are often represented with a feature matrix containing attributes assigned to each node of the graph, and an adjacency matrix – or edge matrix -- describing the connectivity between the nodes. A Graph Neural Networks (GNN) iteratively updates a node’s features integrating the node’s neighborhood information (an operation called message passing). GNNs can be trained to learn the optimal updated node features to predict properties of PPI (Igashov *et al*., 2021; Cao and Shen, 2020; Wang *et al*., 2021). Contrary to CNNs, the convolution operations on graphs can be independent from cartesian coordinates and only rely on the relative local connectivity between nodes, therefore making graph rotational invariant. GNNs are also invariant with respect to the ordering of nodes in the feature and adjacency matrices, and the network can accept any size of graph, therefore more naturally representing the diversity of PPIs.

Building up on our previous framework DeepRank (CNN based), we present here DeepRank-GNN (Reau and Renaud, 2021) that takes advantage of the intrinsic properties of graph representation and graph convolutions. DeepRank-GNN converts interfaces of protein-protein complexes from 3D coordinates PDB files into graphs and enables the application of pre-defined or user-defined GNN architectures to identify problem-specific patterns in PPIs. DeepRank-GNN can automatically compute docking-specific target values when reference PDB files are provided or assign user-provided target values to the graphs. We describe the main functionalities of DeepRank-GNN and showcase its application to the scoring of docking models where we use Graph Interaction Network (GINet) that aims at learning favorable interaction patterns between two subgraphs. Detailed documentation is available online at https://deeprank-gnn.readthedocs.io/.

## 2. Methods

### 2.1. Datasets

#### BM5 Dataset

The Docking Benchmark version 5 (BM5) dataset has been designed for docking purpose and encompasses a non-redundant set of 231 complexes for which the individual structure of interacting proteins is available in a bound and an unbound conformation (Vreven *et al*., 2015). We discarded the 56 antibody-antigen complexes plus the complexes involving more than 2 chains, and worked on the remaining 142 dimers. As described in Renaud Renaud *et al*., 2021, we generated 23 500 models using our integrative modelling software HADDOCK (see SI). The overall dataset comprises 3337000 models and is available from the SBGrid data repository https://data.sbgrid.org/dataset/843/.

#### Training Evaluation and Test sets

We performed two 5-fold cross validation in which the training and the evaluation sets change over the folds while the test set remains constant. The test set consists of all docking models generated for 15 randomly selected complexes (375700 models, 10% of the dataset, see Table S1). Each fold consists of 10% of the docking models per remaining complexes without any models overlap between folds. Their composition preserves the distribution of CAPRI iRMSD classes (Lensink *et al*., 2007) (reporting on the quality of the models) per complex. This was achieved using sklearn StratifiedKFold tool. The 127 complexes not included in the test set are then split into a training (80%, i.e. 102 complexes, 258060 models) and an evaluation set (20%, i.e. 25 complexes, 63250 models per fold). The detailed content of each fold’s training and evaluation set is provided in Table S2-S11. The Graph Interaction Network (GINet) described in the results section was applied using the Markov Clustering Algorithm (MCL) to identify the node clusters used in the pooling operations. A number of complexes displaying important clashes could not be converted into graphs.

#### CAPRI dataset

The CAPRI score set (Lensink and Wodak, 2014a) was used as an external test set. It consists of 13 protein dimers for a total of 16666 models generated by over 40 different research teams using a variety of software. It is acknowledged as the most diverse set of docking models with targets of different complexity.

#### Other scoring functions

The HADDOCK (van Zundert *et al*., 2016), iScore (Geng *et al*., 2020), DOVE (Wang *et al*., 2020) and DeepRank scores were computed on the CAPRI score set as described in Renaud *et al*., 2021. Deeprank and DOVE are two CNN based scoring approaches, iScore is graph-kernel based, and HADDOCK uses a classic scoring function that consists of a linear combination of energy terms (see SI). These scores for the BM5 and CAPRI score sets were obtained from the DeepRank paper (Renaud *et al*., 2021) and can be downloaded from: https://data.sbgrid.org/dataset/843/.

### 2.2. Graph Generation and Target Value Computation

PPIs were converted into graph using the default DeepRank-GNN parameters, default nodes features (residue type, polarity, charge and BSA) and edge feature (distance) and additional PSSM information (PSSM profile, PSSM information content, PSSM conservation score). By providing the reference structure of the complex, the bound conformation, DeepRank-GNN automatically computed the fraction of native contacts (*f_nat_*) that was used as the target value (see Table S12). As compared to the interface RMSD (iRMSD) or ligand RMSD (lRMSD) values, the *f_nat_* value is caped between 0 and 1, thus giving the same weight to all bad quality model. For instance, two models very distant from the reference structure will be assigned a 0 *f_nat_* value while they can be assigned 2 very distinct iRMSD values (Fig. S1), which can uselessly influence the network parameters optimization.

### 2.3. Training

The network was trained over 20 epochs on batches of 128 shuffled graphs. We used the mean square error loss (MSE loss) function using the *f_nat_* values as the ground truth and the Adam algorithm (Kingma and Ba, 2017) with a learning rate of 0.001 to minimize the loss. A complete epoch (23500 3D models) required 2.4±0.9 hours on 1 GPU (GeForce GTX 1080 Ti).

### 2.4. Metrics Computation

The Area Under the ROC Curve (AUC), the hit rate and the success rate are computed to evaluate the performance of the scoring functions. To meet the requirement of these metrics that evaluate the discriminating ability of a binary classifier, we binarized the *f_nat_* data using a 0.3 threshold: docking models with a *f_nat_* ≥ 0.3 are considered to be of acceptable quality, while those with a *f_nat_* < 0.3 are considered non-acceptable. This threshold differs from the CAPRI standard *f_nat_* “acceptable” quality threshold of 0.1 that is combined with additional iRMSD and lRMSD criteria (Lensink *et al*., 2007). Herein, since no RMSD values are considered, we raised the acceptance threshold to the equivalent of the CAPRI standard *f_nat_* “medium” quality threshold of 0.3 to avoid misclassifying poor quality models (Fig S1).

The ROC curve is defined as the fraction of True Positive Rate as a function of the fraction False Positive Rate while navigating through the ranking provided by the scoring function. The AUC is the integral of the ROC curve and is equal to 1 for an ideal classifier and 0.5 for a random classifier. The hitrate is defined as the percentage of hits retrieved within the top N ranks. The success rate is the number of complexes for which at least one acceptable quality model is retrieved within the top N

## 3. Results

### 3.1. DeepRank-GNN Overview

DeepRank-GNN is a python3 package based on PyTorch (Paszke *et al*., 2019) that offers a complete framework to learn PPI patterns in an end-to-end fashion using Graph Neural Networks. The DeepRank-GNN framework consists of two mains parts (Fig. 1) the conversion of 3D protein-protein complexes into interaction graphs with node and edge features, 2) the training and the evaluation of a graph neural network model. An overview of the software architecture is provided in Fig. S2. We briefly present both parts below and refer the reader to the online documentation for further information (https://deeprank-gnn.readthedocs.io/).

**Fig. 1.**
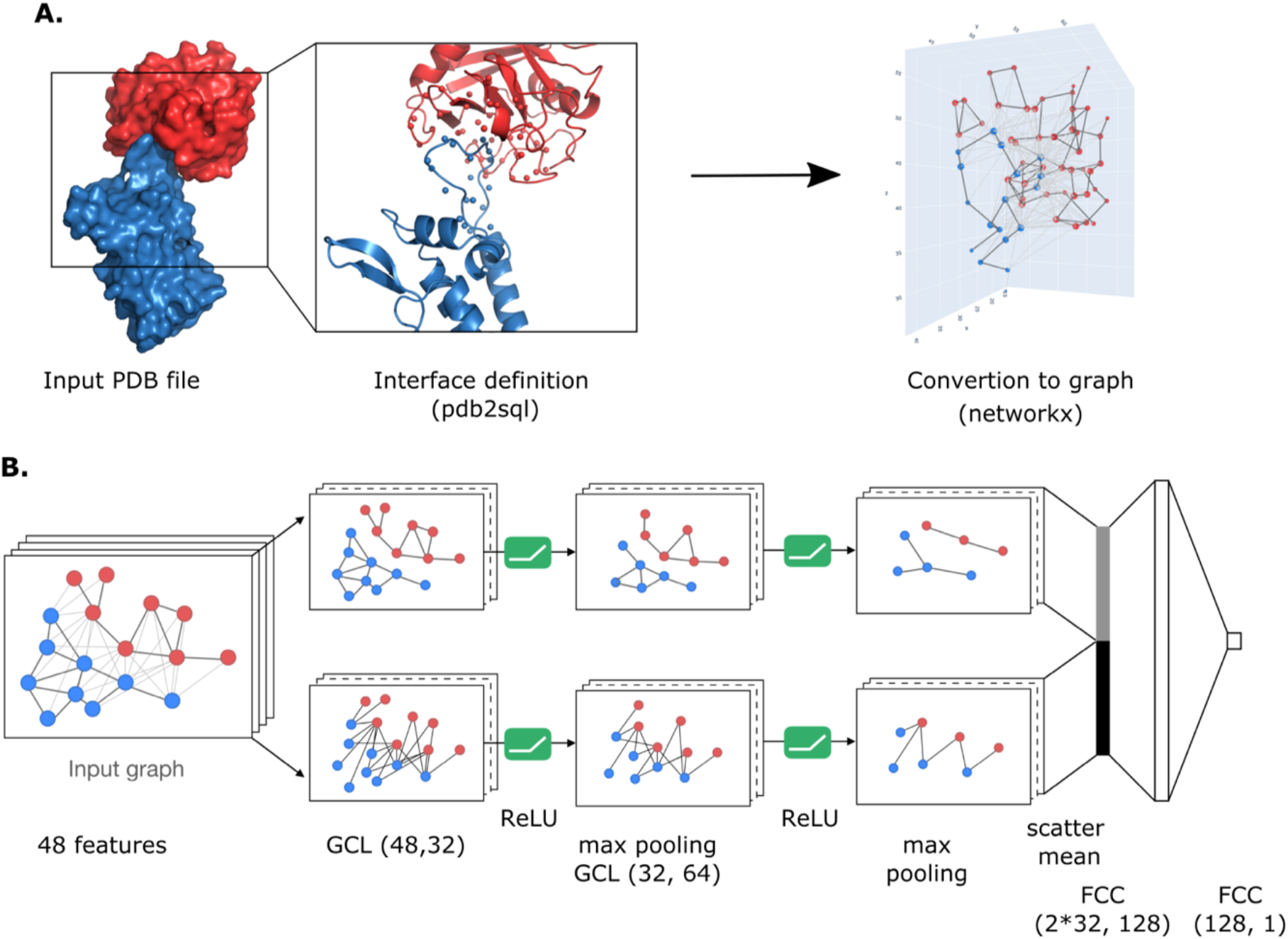
Overview of the DeepRank-GNN framework. (**A**) DeepRank-GNN identifies interface residues and converts them into an interface graph. Internal edges are defined between residues from the same chain having heavy atoms within a 3 Å distance cutoff from each other, while external edges are defined between residues from different chains having heavy atoms within the 8.5 Å cutoff. (B) Architecture of a Graph Interaction Network (GINet). The graph representation of a PPI is split into 2 sub-graphs, i.e. the internal graph connecting atoms from the same protein and the external graph connecting atoms from distinct proteins. The 2 sub-graphs are sequentially passed to 2 consecutive convolution/activation/pooling layers. The two final graph representations are flattened using a scatter mean operation and merged before applying two fully connected layers. GCL: graph convolution layer; FCC: fully connected layer.

#### Graph Generation

DeepRank-GNN converts protein-protein complexes into residue level-graph that are focused on the interface between the proteins (Fig. 1.A.). It takes PDB 3D coordinate files as an input and defines the interface between two chains using pdb2sql (Renaud and Geng, 2021b), our PDB file parser using Structured Query Language (SQL). By default, the interface is defined by all the residues involved in intermolecular contacts, i.e. the residues of a given chain having a heavy atom within an 8.5 Å distance cutoff of any heavy atom from another chain. These contact residues form the nodes of the graph. *Interface* edges are defined between two contact residues from distinct chains presenting a minimal atomic distance smaller than 8.5 Å. In addition, *internal* edges are defined between two contact residues of the same chain provided they have heavy atoms within 3 Å from each other. These distance cut-offs can be tailored by the user. The types of edges can be later considered to perform different convolution operations on the graph.

Graphs are stored in HDF5 format that is suited for large dataset storage and allows efficient memory usage and fast input/output operations during the network training.

#### Featurization

By default, DeepRank-GNN computes and assigns an ensemble of residue-level features to each node. Those are summarized in Table 1. The feature computation can rapidly become a limiting step if the computation speed is not optimized. In DeepRank-GNN, the assignment of residue type, charge, polarity and buried surface area features are considerably faster than the computation of the residue depth, i.e. the average distance of the atoms of a residue from the solvent accessible surface, and the half sphere exposure. Provided that the information brought by the two latter could implicitly be deduced from the buried surface area feature and the node environment, they are not calculated by default for the sake of time efficiency. Pre-computed Position Specific Scoring Matrices (PSSM) matrices are required for the computation of PSSM related features. We strongly advise querying dataset of precomputed PSSM matrices such as the 3DCONS (http://3dcons.cnb.csic.es/), the Conserved Domains Database (Lu *et al*., 2020) or using our in-house PSSM generation tool PSSMGen (Renaud and Geng, 2021a).

**Table 1.** Features computed in DeepRank-GNN.

| Name of the feature | Full name | Description | Default | Number of parameters | Type |
| --- | --- | --- | --- | --- | --- |
| type | Residue Type | One-hot encoded | Default | 20 | Node feature |
| charge | Residue Charge |  | Default | 1 | Node feature |
| polarity | Residue Polarity | One-hot encoded | Default | 4 | Node feature |
| BSA | Buried Surface Area | FreeSASA | Default | 1 | Node feature |
| PSSM | Position-Specific Scoring Matrix |  | Optional | 20 | Node feature |
| cons | Conservation Score – from PSSM |  | Optional | 1 | Node feature |
| ic | Information Content – from PSSM |  | Optional | 1 | Node feature |
| depth | Residue Depth | MSMS - Biopython | Optional | 1 | Node feature |
| hse | Residue Half sphere exposure | Biopython | Optional | 1 | Node feature |
| distance | Normalized distance |  | Default | 1 | Edge feature |

To encode the relative positions of the nodes in the graph, and therefore the overall structure of the interface, we assign a distance feature to the internal and external edges. This distance feature is based on the smallest atomic distance between 2 residues (nodes) that is transformed into an interaction strength by equation 2.

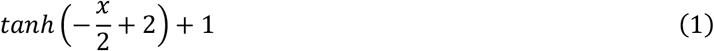

The interaction strength ranges from 0 for long distances to 2 for short distances, and provides a normalized feature of the internode distance.

#### Target assignment

Many different metrics have been developed to quantify the relevance of PPIs. In a docking scenario where the goal is to identify near-native models, the user can provide a reference structure for DeepRank-GNN to automatically compute target values based on CAPRI quality criteria (Lensink *et al*., 2007) (Table S12). For other applications and/or use cases, users may input their own problem-specific target values or develop new metric calculations and integrate these metrics in the computational workflow.

#### Model Training/Evaluation/Test

All the prerequisite to train and to evaluate a GNN model are detailed in our online documentation. The user can run DeepRank-GNN in a regression or classification mode. The loss function will be automatically set to Mean Square Error (MSE) for regression tasks or to cross-entropy for classification tasks. Weights can be assigned to classes to balance the cross-entropy loss calculation in case of unbalanced dataset. We also propose an automated weight computation that assigns weights inversely proportional to each class representation in the training set for classification tasks on imbalanced datasets (see SI).

#### Network

DeepRank-GNN provides a flexible structure allowing users to define their own network architectures or use predefined ones. We introduce here a new GNN architecture, dubbed Graph Interaction Network (GINet), whose general structure is represented in Fig. 1.B. As seen in this figure GINets are composed of a succession of graph convolution layers (GCL), non-linear activation (here ReLU) and pooling layers. Two distinct GCLs are applied at each convolution steps. One GCL is applied on interface graphs, i.e. graphs with edges connecting nodes from distinct proteins, and a distinct GCL is used on internal graphs, i.e. graphs with edges connecting nodes from the same protein. The rationale behind this architecture is to extract information not only on the interaction itself but also on the propensity of each individual interface to establish an interaction.

Following the convolution/activation/pooling layers, graphs are flattened using the mean value of each node feature and the new feature tensors are concatenated and fed to two fully connected layers with an intermediate dropout layer (with p=0.4).

#### GINet Convolution Layers

The convolution layers in the GINet are inspired by the graph attention network (GAT) described by Veličković *et al*., 2018 and the edge aggregated graph attention network (EGAT) from Mahbub and Bayzid, 2020. Attention mechanism (Veličković *et al*., 2018) is used to weight the contribution of individual neighbors in the new state of a node. When applied to PPIs, we except attention mechanism to learn favorable or deleterious contacts in a local environment.

The feature representation of node i, 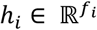, is transformed into a new feature representation 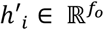 (where *f_i_* and *f_o_* are respectively the number of input and output features) through the aggregation of the feature representation of neighbouring nodes 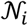, i.e. the first ordered neighbours of i, including node i itself. We use the weighted sum of the neighbours feature representation as an aggregator,

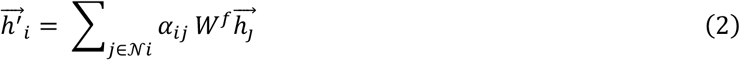

where 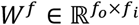 is the learnable parameters used in the linear transformation of the node features representation and where the weights *α_ij_*are learned through an attention mechanism described in equations 4 and 5. Each edge of the graph is assigned an attention score *s_ij_* given by

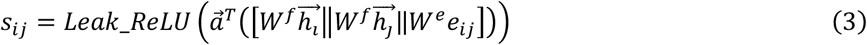

Where *e_ij_* is edge feature, 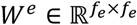 is the learnable parameter used in the linear transformation of the edge feature representation (*f_e_* being the number of edge features), and 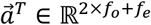 is the attention mechanism. || denotes vector concatenation. *s_ij_* is further normalized by a SoftMax activation function to give a probability distribution *α_i_* (between 0 and 1) over the node neighbours 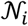, with *α_ij_* reflecting how much the neighbouring node j should contribute to the new representation of node i.

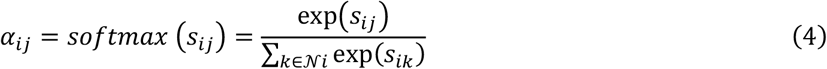

As described by Mahbub and Bayzid, 2020 and shown in equation 4, the calculation of the attention score considers the node features 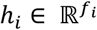 and 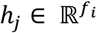 as in the original GAT paper plus the edge feature 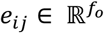. The addition of the edge feature, which herein corresponds to the interaction strength, is essential for PPIs studies as it is well established that the potential energy of a system depends on the atomic pairwise distances. In a coarse-grained system with residue-level representation, the same trend transfers to residue-residue distances.

#### Clustering and Pooling Layers

Pooling layers are used to reduce the number of nodes in the graph, which depends on the input 3D model, and learn higher level features, i.e. features of group of nodes, during the training. The pooling operation is here performed on clusters of highly interacting nodes that can be computed using either a Markov Cluster Algorithm (MCL)(Enright *et al*., 2002), or the Louvain community detection algorithm (Blondel *et al*., 2008). Both clustering methods rely on the interaction strength and therefore cluster nodes that are close from each other in 3D space on the flight. A max pooling is then applied on these clusters. This pooling operation aggregates the nodes of a given cluster in a single node whose feature values are given by the maximum feature values across the cluster. Interface and internal edges are then drawn to connect these new nodes. The edge features are here obtained by summing up all the edge features of the nodes belonging to the cluster.

#### Quality Metrics

DeepRank-GNN provides tools to swiftly compute the quality metrics summarized in Table 2. Upon the definition of a threshold value to binarize the data, all classification metrics can be applied to continuous targets and prediction values.

**Table 2.** Prediction performance quality metrics computed in DeepRank-GNN.

| Classification |  |
| --- | --- |
| sensitivity | Sensitivity, hit rate, recall, or true positive rate |
| specificity | Specificity or true negative rate |
| specificity | Precision or positive predictive value |
| NVP | Negative predictive value |
| FPR | Fall out or false positive rate |
| FNR | False negative rate |
| FDR | False discovery rate |
| accuracy | Accuracy |
| Regression |  |
| Explained_variance | Explained variance score function |
| max_error | Maximum residual error |
| mean_absolute_error | Mean absolute error loss |
| mean_squared_error | Mean squared error loss |
| root_mean_squared_error | Root mean squared error loss |
| mean_squared_log_error | Mean squared logarithmic error loss |
| median_squared_log_error | Median absolute error loss |
| r2_score | R<sup>2</sup> coefficient of determination |

### 3.2. Ranking of PPI docking models

Docking is an *in silico* modelling approach commonly used to predict the 3D structure of biomolecular complexes. Docking involves two steps: The sampling, i.e. the exploration of the conformational interaction space to generate 3D models, and the scoring that aims to identify near-native models out of the pool of generated docking models. As illustrated by the Critical Assessment of PRedicted Interactions (CAPRI) initiative that frequently proposes blind predictions of experimentally determined 3D structures of protein complexes, there is still room for scoring functions improvement (Lensink *et al*., 2016; Lensink *et al*., 2021). Most scoring functions can be classified into physical energy-based, statistical potential-based and machine learning based functions. They are constantly explored for improvement to either propose system-specific or broad-spectra scoring tools, which in some cases are also used to predict changes in binding affinities (Geng *et al*., 2019).

Here, we demonstrate the use DeepRank-GNN to score docking models of various complexes generated with a variety of docking software.

#### Performance of 10-fold cross validation on the BM5

Ten-fold cross validation was performed to analyze the performance and robustness of DeepRank-GNN on the task of scoring docking models. We trained here our model on the BM5 data set using 258 060 structures from 102 distinct complexes (see Methods). The trained GINet models were then validated on 63 250 structure from 25 complexes (see Fig. S3). The training and validation set were distinct at the complex level, i.e. the trained network has never been trained on any structure from the complexes used for validation. As described in the method section, the training and the evaluation set change over the folds while the test set remains constant. For each fold, we retained the generated model that minimizes the most the loss value on the evaluation set and evaluated its performance on the BM5 test set that consists of 375700 structures from 15 complexes (described in Table S1). As shown in Fig. 2A, when defining positive docking models as those associated to a *f_nat_* ≥ 0.3 and averaging the true positive rate (TPR) over the number of BM5 test complexes, most GINet models globally perform equally or better than the HADDOCK scoring function (see SI) for 8 out of 10 DeepRank-GNN models, yielding an AUC ≥ 0.95 on the test set. Nine DeepRank-GNN models out of 10 display a lower false positive rate (FPR) (0.03 ≤ FPR ≤ 0.07) as compared to HADDOCK (0.05 ≤ FPR ≤ 0.07) for TPRs between 0.62 and 0.72. We however notice a variation in the performance depending on the data subset used for the training and the evaluation of the models, which is particularly clear when we consider not only the complex-averaged TPR, but also the standard deviation (Fig. S4) and the hit rates obtained on individual complexes (Fig. S5, highlighting the dataset dependency of DeepRank-GNN performance. Among all DeepRank-GNN models, highest performance is reached with the one generated in the fold6 (AUC = 0.97±0.3).

**Fig. 2.**
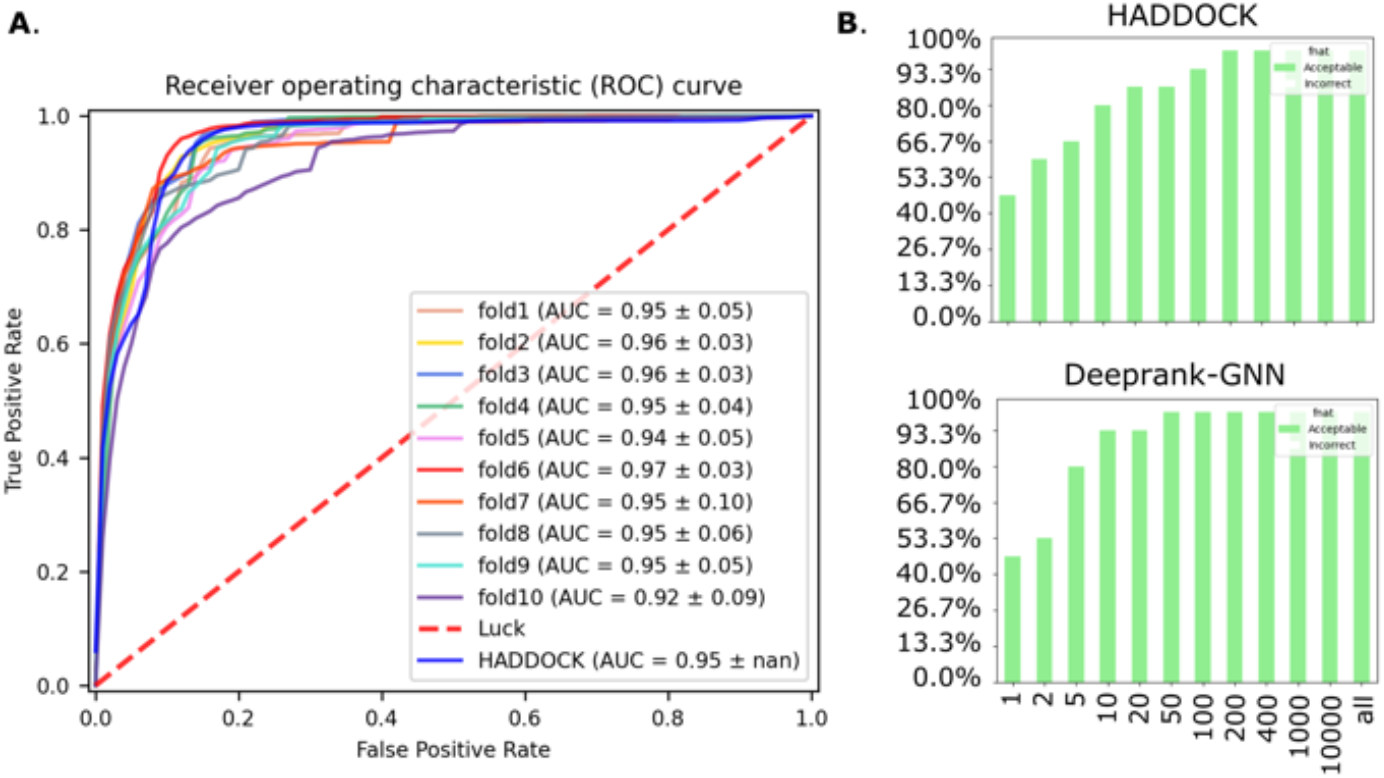
Comparison of DeepRank-GNN with HADDOCK scoring function on the BM5 set. (A) Average ROC curves obtained with the models retained for each DeepRank-GNN fold and HADDOCK score. A true positive case corresponds to a complex with fnat > 0.3 correctly predicted. The number of True Positive Rate value is averaged over the number of complexes in the test dataset. The dashed line represents a random classifier. (B) Success rates of HADDOCK and DeepnRank-GNN on the BM5 test dataset.

#### Rank Correlation

Since rank correlation is a good indicator of the predictiveness of a score, we computed the spearman *ρ* correlation between the *f_nat_* and the DeepRank-GNN scores obtained with the best GINet model (i.e. the fold6 model) on the entire test set (it0/it1/itw). We observe an average Spearman correlation of 0.5±0.13, the highest correlation being obtained for 1PPE (*ρ* = 0.64), the lowest for 1IBR (*ρ* = 0.22) (Fig. S6). Interestingly, 1PPE constitutes an *easy* case with 11.5% of good docking models (*f_nat_* ≥ 0.3) generated, while 1IBR constitute a difficult case with 0.4%. This 0.4% correspond to the 100 models obtained by refinement of the bound complex and display very low structural diversity. Overall, we observe a good ability to identify near-native models in the top-ranked model with impressive success rates of 47, 80 and 93% (7, 12 and 14 over 15 test complexes) at top1, top5 and top10 respectively.

The performance considerably increases when considering only the it1/itw models of the test set with an average Spearman correlation of 0.68±0.18, the highest correlation being obtained for 2SNI (*ρ* = 0.86), the lowest for 1F6M (*ρ* = 0.22) (Fig. S7). Here again, 1F6M constitutes a difficult case with 102 good models in the entire pool of docking models, 100 of them consisting in refined structure of the bound complex and representing very low diversity in the binding mode. The success rate is similar to the one on the entire set with values of 53, 87 and 93% (8, 13 and 14 over 15 test complexes) at top1, top5 and top10 respectively. In both scenario, success rates of 47, 67 and 80% (7, 10 and 13 over 15) are obtained with HADDOCK scores for the top1, top5 and top10, respectively (Fig. 2.B).

#### Performance on the external CAPRI Score test set

We evaluated the performance of DeepRank-GNN (fold6 model), DeepRank, DOVE, HADDOCK and iScore on the CAPRI score set. DeepRank and DOVE are two CNN based scoring approaches, iScore is graph-kernel based, and HADDOCK uses a classic scoring function that consists of a linear combination of energy terms. The CAPRI score set consists of 13 complexes for which 497 to 1987 models have been generated by different groups using a wide diversity of docking tools and protocols(Lensink and Wodak, 2014b). When considering the AUC DeepRank-GNN stands on top together with iScore with an average AUC of 0.68 and 0.64 respectively (Fig. 3.A). However, in terms of early enrichments (success rate of top N models for N≤10) iScore scores best followed by DeepRank-GNN and HADDOCK (Fig. 3.B). Note that both iScore and DeepRank-GNN are using graph representations and both encode PSSM as a key feature.

**Fig. 3.**
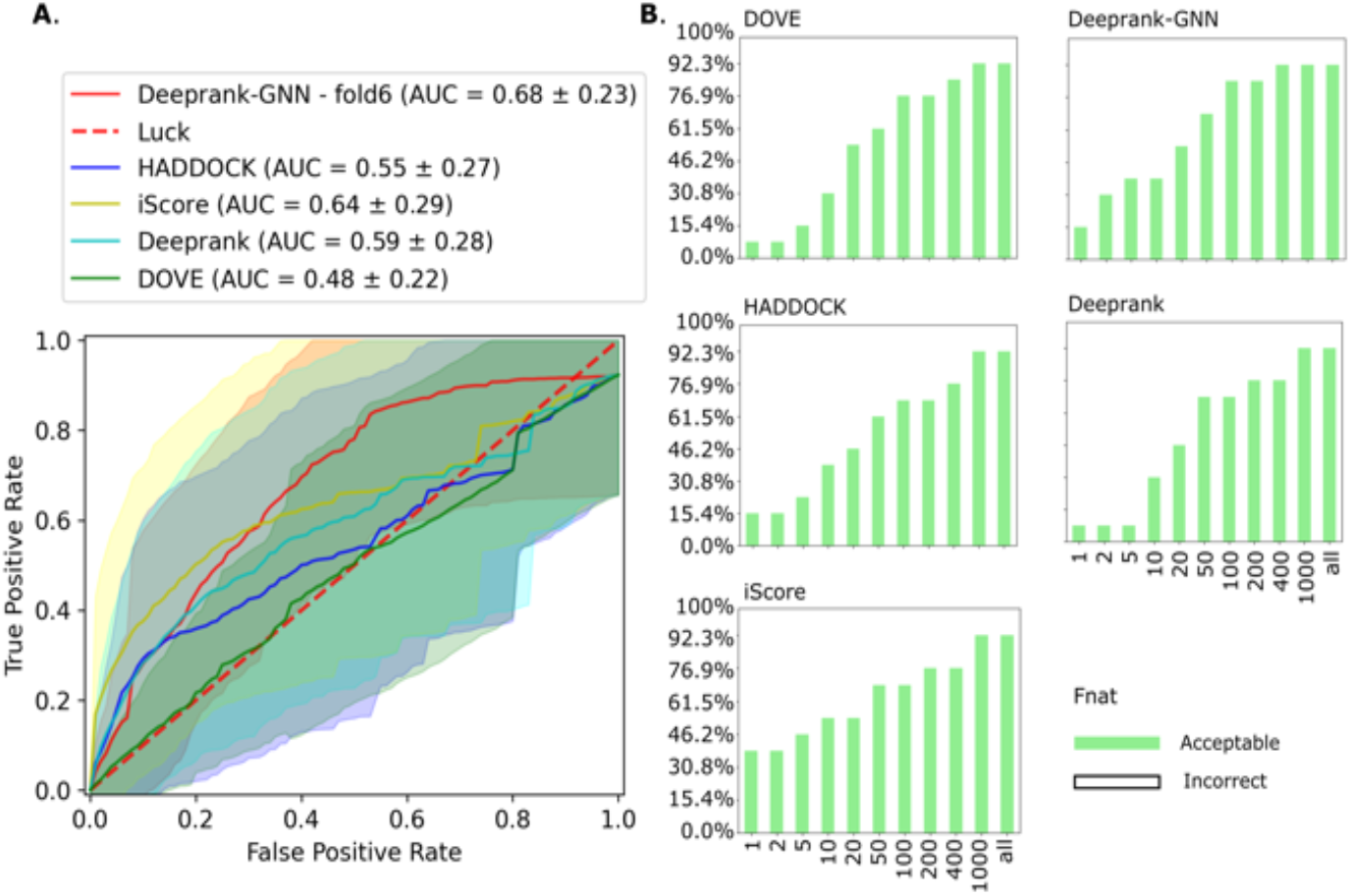
Comparison of the performance obtained on the CAPRI Scoreset. See legend of Fig.2.

### 3.3. Computational performance

The graph representation of the interface not only provides a natural way of representing PPIs, it also considerably improves the computation performance in terms of storage, data generation and learning speed as compared to the use of grids and CNN. To quantify it, we compared the graph generation step of DeepRank-GNN to the grid generation step of DeepRank on the CAPRI score set (Table S13-15) as well as each protocol’s training speed (Table S16) using MPI distributed processes on 4 CPUs. The results show that the graph generation is on average 20 times faster than the 3D grid generation in CNN (0.65 ± 0.31 vs 12.4 ± 3.3 second per model) and requires ~22 times less storage space (0.14 ± 0.1 vs 3.07 ± 0.4 MB per model). It is worth noting that default settings were used for each approach and that DeepRank computes additional atomic-level descriptors leading to a total of 72 descriptors against 48 in DeepRank-GNN (Table S17). To fairly compare the computation performance in the training phase, we retained comparable residue-level features for the two protocols corresponding to 48 and 58 features for DeepRank-GNN and DeepRank respectively, and split the CAPRI score set into 80% training and 20% evaluation sets. We trained the GINet model of DeepRank-GNN described in section 3.1 and the default 3D-CNN implemented in DeepRank over 10 epochs. The results show a remarkable speed difference, DeepRank-GNN being ~25 times faster than DeepRank when no data augmentation is used in the latter (Table S16).

## 4. Conclusion

We have developed DeepRank-GNN, a new computational framework to learn and predict interaction patterns from protein-protein interfaces (PPI). DeepRank-GNN is provided a freely accessible python3 package (https://github.com/DeepRank/DeepRank-GNN). The framework encompasses pre-processing tools that take PPI PDB files as input, converts the interface of interaction into residue-level graphs and automatically assigns biologically relevant features to the graphs. In a second step, the generated graphs can be used to train, evaluate and test a provided or user-defined GNN to make problem specific predictions. DeepRank-GNN has been designed to be applied to various PPI-related project and offers the user many options such as the possibility to select appropriate features, to use any type of target values, to reweight the scoring functions for classification tasks, to design his/her own GNN architecture etc.

As a demonstration we applied DeepRank-GNN to the task of scoring docking models of various complexes from the Docking Benchmark 5 (BM5) and the CAPRI score set. From the 147 selected BM5 complexes, 90% were used to train the model by performing a 10-fold cross validation. The remaining 10% constitute the BM5 test set. The trained models (one per fold) globally display a data dependency, yet most of them compete with- or outperform the HADDOCK scoring function. The scoring performance of the best model, namely fold6-model, was further evaluated on the CAPRI score set and compared to HADDOCK, DeepRank, DOVE and iScore. iScore and DeepRank-GNN rank 1^st^ and 2^nd^ in this task. Interestingly, iScore and DeepRank-GNN approaches lie on a simplified residue-level graph representation of the surface of interaction between the two proteins of the complex, while other approaches deal with atom-level representation of the complex. They also both account for conservation information by the mean of PSSM features. If this information is likely to contribute to such performance, it is however not sufficient to explain the better reached performance as compared to the other explored tools provided that DeepRank also considers PSSM and displays poor early enrichment on the CAPRI score set. Additionally, among all scoring function mentioned in this work, DeepRank-GNN is the only one that does not contain energy terms, highlighting that simple geometric and physico-chemical properties could be as informative as (or more than) the approximative energy terms used in most scoring functions. Further optimization of the hyperparameters of the GINet or the design and application of new GNNs could further improve the performance observed here.

Overall, this framework can be used for various other applications (e.g. binding affinity and physiological interface prediction) and offers many perspectives of extension such as the application to single proteins (e.g. for genetic variant pathogenicity prediction), to larger multimeric states (e.g.for larger complex quality prediction), or the integration of other GNN architectures such as the E(n)-equivariant graph neural networks (Satorras *et al*.,2021) to run molecular dynamics simulations.

## Software availability

The software is available on Github (https://github.com/DeepRank/Deeprank-GNN) and the documentation is available online (https://deeprank-gnn.readthedocs.io/).

## Data availability

The BM5 and CAPRI score set docking models are obtained from the DeepRank paper (Renaud *et al*., 2021) and can be downloaded from: https://data.sbgrid.org/dataset/843/.

## Funding

The project was supported by an ASDI grant from the Netherlands eScience Center (grant number ASDI.2016.043), by a SURF Open Lab “Machine learning enhanced HPC applications” grant (AB/AM/10573), by a “Computing Time on National Computer Facilities” grant (2018/ENW/00485366) from the Netherlands Organization for Scientific Research (NWO). MR and AMJJB acknowledge financial support from the European Union Horizon 2020 project BioExcel (823830) and from a TOP-PUNT grant (718.015.001).

### Conflict of Interest

none declared.

## Notes

https://github.com/DeepRank/DeepRank-GNN

https://deeprank-gnn.readthedocs.io/

https://data.sbgrid.org/dataset/843/

### Competing Interest Statement

The authors have declared no competing interest.

