## Supplementary Information for "DeepRank-GNN: A Graph Neural Network Framework to Learn Patterns in Protein-Protein Interfaces"

*To whom correspondence should be addressed.

\# These authors contributed equally.

### Table of content

### Tables and Figures

### Generation of docking models with HADDOCK

Docking with HADDOCK follows a three step process: 1) in the first stage (it0), the docking is performed from the separated and randomly rotated starting conformations, treating the proteins as rigid units,; 2) in the second stage (it1), a semi-flexible refinement is performed, consisting of a simulated annealing in torsional space (with fixed bond lengths and angles) introducing flexibility first along the side chains at the interface (i.e. those within 5 Å of the partner protein), and second in both backbone and side chains of the interface residues; 3) in the last stage (itw), the models are subjected to a final energy minimization (and/or short refinement in explicit solvent (water)). Each HADDOCK step has its own scoring function and restraints can be provided to guide the docking with a priori information.

- ${HADDOCK}_{it0}=0.01\times E_{vdw}+1.0 \times E_{elec}+1.0 \times E_{desolv}-0.01 \times BSA$
- ${HADDOCK}_{it1}=1.0\times E_{vdw}+1.0 \times E_{elec}+1.0 \times E_{desolv}-0.01 \times BSA$
- ${HADDOCK}_{itw}=1.0\times E_{vdw}+0.2 \times E_{elec}+1.0 \times E_{desolv}$

In order to ensure a suitable number of near-native models (i.e. iRMSD to the reference structure ≤ 4 Å), we combined models from 5 docking scenarios with increasing level of a priori information: 1) docking with random surface patch restraints (10000/400/400 models for it0/it1/water stages), 2) docking with center of mass restrains (10000/400/400 models for it0/it1/water stages), 3) docking with true interface residues defined within a 5 A distance to the partner protein (1000/400/400 for it0/it1/water stages), 4) docking with true interface residues defined within a 3.9 A distance to the partner protein (1000/400/400 for it0/it1/water stages) and 5) refinement of the bound complex (50/50 it1/water stages).

### Automated weights computation for CrossEntropy Loss function

The cross-entropy loss function is commonly used for classification tasks and more especially when dealing with multiple classes. The cross-entropy loss function computes the probability of each class y, and applies a logarithmic penalty based on how far is the prediction from the ground truth value.

For a binary classification tasks, the cross-entropy loss function is defined as:

$${Loss}_{CE}=-\sum_{i=1}^{2} y_{i}\log\left( p_{i} \right)= -(y\log\left( p \right)+\left( 1-y \right)\log\left( 1-p \right))$$

Where i spans the number of classes, $y_{i}$ is the ground truth value (0 or 1), and $p_{i}$ is the probability of the class i, and where $p_{1}=1- p_{2}$. Provided that $y_{n}$is the target (i.e. the correct class to predict), then $y_{n}=1$ and all other $y_{i}=0$. We can thus simplify the equation for multi-classes tasks to:

$${Loss}_{CE}=-y_{n}\log\left( p_{n} \right)$$

$p_{n}$being the probability of the target class $y_{n}$ and being computed with the SoftMax activation function:

$${Loss}_{CE}=-y_{n}\log\left( \frac{exp(x_{n,y_{n}})}{\sum_{c=1}^{C} exp(x_{n,c})} \right)$$

where $x_{n,y_{n}}$ is the predicted value for the target class, c spans the number of classes, and $x_{n,c}$ is the predicted value for each class c.

In pytorch, the cross-entropy loss is computed for each entry n of the input batch as follow:

$${Loss}_{CE}=\left\{ l_{1},l_{2},\ldots,l_{N} \right\};$$

With

$$l_{n}= -{w_{y_{n}}y}_{n}\log\left( \frac{exp(x_{n,y_{n}})}{\sum_{c=1}^{C} exp(x_{n,c})} \right)= -w_{y_{n}}\log\left( \frac{exp(x_{n,y_{n}})}{\sum_{c=1}^{C} exp(x_{n,c})} \right)= -w_{y_{n}}\log(p_{n})$$

$w_{y_{n}}$being the weight, or scaling factor, assigned to the target class. By default, $w_{y_{n}}$ is set to 1 for each class c, meaning that all classes equally contribute to the loss calculation. The batch loss can be computed as the mean (default in pytorch and DeepRank-GNN) or the sum of individual losses. However, in case of unbalanced training dataset, it is recommended to weight the different classes with higher weights for the minority class(es) to avoid optimizing the neural network weights solely based on majority classes. An agreed upon technique to weight the contribution of each classes in the loss function is to assign weights that are inversely proportional to the frequency of each class in the training set.

In DeepRank-GNN, the users can input their own weights or let DeepRank-GNN automatically compute them as follow:

- Compute the frequency of each class
- Convert it into a percentage

Example :

We have a training dataset of 400 graphs, 4 classes (0,1,2,3) split as follow:

- Class 0: 200 graphs
- Class 1: 50 graphs
- Class 2: 50 graphs
- Class 3: 100 graphs

The frequency F is given by:

$F=\left[ \frac{1}{200}, \frac{1}{50},\frac{1}{50},\frac{1}{100} \right]= [0.005, 0.02, 0.02, 0.01]$

It is then transformed into a frequency percentage:

$weights=\left[ \frac{0.005}{\sum_{i=0}^{3} F_{i}}, \frac{0.02}{\sum_{i=0}^{3} F_{i}},\frac{0.02}{\sum_{i=0}^{3} F_{i}},\frac{0.01}{\sum_{i=0}^{3} F_{i}} \right]= [0.0909, 0.3636, 0.3636, 0.1818]$

Table S1 Composition of the test set.

|  | **fnat >= 0.3** | **fnat < 0.3** | **total** | **fraction good models** | DeepRank-GNN scoring time per complex  (seconds) | DeepRank-GNN scoring time per model  (seconds) |
| --- | --- | --- | --- | --- | --- | --- |
| **1AK4** | 2352 | 22948 | 25300 | 9,3 % | 900,7 | 3,6E-02 |
| **1BVN** | 2337 | 22963 | 25300 | 9,2 % | 861,0 | 3,4E-02 |
| **1CGI** | 751 | 24549 | 25300 | 3,0 % | 946,0 | 3,7E-02 |
| **1F6M** | 102 | 25197 | 25299 | 0,4 % | 874,8 | 3,5E-02 |
| **1H1V** | 211 | 11610 | 11821 | 1,8 % | 228,5 | 1,9E-02 |
| **1IBR** | 100 | 25198 | 25298 | 0,4 % | 970,5 | 3,8E-02 |
| **1OPH** | 100 | 25199 | 25299 | 0,4 % | 886,0 | 3,5E-02 |
| **1OYV** | 1951 | 23349 | 25300 | 7,7 % | 457,4 | 1,8E-02 |
| **1PPE** | 2913 | 22386 | 25299 | 11,5 % | 467,3 | 1,8E-02 |
| **1XQS** | 2091 | 23209 | 25300 | 8,3 % | 449,5 | 1,8E-02 |
| **2OZA** | 693 | 22807 | 23500 | 2,9 % | 426,4 | 1,8E-02 |
| **2SNI** | 2488 | 22812 | 25300 | 9,8 % | 465,7 | 1,8E-02 |
| **2YVJ** | 1215 | 24085 | 25300 | 4,8 % | 704,9 | 2,8E-02 |
| **2Z0E** | 257 | 25043 | 25300 | 1,0 % | 823,1 | 3,3E-02 |
| **3K75** | 3293 | 21986 | 25279 | 13,0 % | 877,1 | 3,5E-02 |
|  |  |  | average % of good models | 5,6 % | average speed per model | 2,8E-02 |
|  |  |  | standard deviation | 4,5 % |  |  |

Table S2 Training and evaluation set composition – fold1

| fold1 | fnat ≥ 0.3 | fnat < 0.3 | set | total | % good models |
| --- | --- | --- | --- | --- | --- |
| 1ACB | 5 | 1339 | train | 1344 | 0,4 % |
| 1ATN | 28 | 2021 | train | 2049 | 1,4 % |
| 1AVX | 159 | 1592 | train | 1751 | 9,1 % |
| 1AY7 | 221 | 2309 | train | 2530 | 8,7 % |
| 1B6C | 82 | 2448 | train | 2530 | 3,2 % |
| 1BKD | 12 | 1789 | train | 1801 | 0,7 % |
| 1BUH | 203 | 2324 | train | 2527 | 8,0 % |
| 1CLV | 15 | 2515 | train | 2530 | 0,6 % |
| 1D6R | 79 | 2451 | train | 2530 | 3,1 % |
| 1DFJ | 24 | 2414 | train | 2438 | 1,0 % |
| 1E+96 | 269 | 2257 | train | 2526 | 10,6 % |
| 1E6E | 253 | 2276 | train | 2529 | 10,0 % |
| 1EAW | 100 | 2425 | train | 2525 | 4,0 % |
| 1EFN | 126 | 2404 | train | 2530 | 5,0 % |
| 1EWY | 146 | 2384 | train | 2530 | 5,8 % |
| 1F34 | 80 | 2443 | train | 2523 | 3,2 % |
| 1FC2 | 94 | 2427 | train | 2521 | 3,7 % |
| 1FFW | 129 | 2401 | train | 2530 | 5,1 % |
| 1FLE | 88 | 2442 | train | 2530 | 3,5 % |
| 1FQ1 | 112 | 2418 | train | 2530 | 4,4 % |
| 1FQJ | 143 | 2387 | train | 2530 | 5,7 % |
| 1GCQ | 261 | 2269 | train | 2530 | 10,3 % |
| 1GHQ | 118 | 2412 | train | 2530 | 4,7 % |
| 1GL1 | 50 | 2480 | train | 2530 | 2,0 % |
| 1GLA | 25 | 2505 | train | 2530 | 1,0 % |
| 1GPW | 50 | 2480 | train | 2530 | 2,0 % |
| 1GRN | 169 | 451 | eval | 620 | 27,3 % |
| 1GXD | 213 | 2317 | train | 2530 | 8,4 % |
| 1H9D | 37 | 2487 | train | 2524 | 1,5 % |
| 1HE1 | 137 | 2023 | eval | 2160 | 6,3 % |
| 1HE8 | 85 | 2445 | eval | 2530 | 3,4 % |
| 1I2M | 11 | 2519 | train | 2530 | 0,4 % |
| 1J2J | 257 | 2273 | train | 2530 | 10,2 % |
| 1JIW | 251 | 839 | eval | 1090 | 23,0 % |
| 1JK9 | 240 | 2290 | eval | 2530 | 9,5 % |
| 1JTD | 125 | 2346 | train | 2471 | 5,1 % |
| 1JTG | 101 | 2429 | train | 2530 | 4,0 % |
| 1KAC | 164 | 2366 | train | 2530 | 6,5 % |
| 1KTZ | 122 | 2408 | train | 2530 | 4,8 % |
| 1KXP | 162 | 2368 | train | 2530 | 6,4 % |
| 1KXQ | 250 | 1692 | train | 1942 | 12,9 % |
| 1LFD | 55 | 2475 | train | 2530 | 2,2 % |
| 1M10 | 10 | 2520 | train | 2530 | 0,4 % |
| 1MAH | 294 | 2236 | train | 2530 | 11,6 % |
| 1MQ8 | 277 | 2253 | eval | 2530 | 10,9 % |
| 1NW9 | 38 | 2492 | eval | 2530 | 1,5 % |
| 1OC0 | 67 | 1013 | train | 1080 | 6,2 % |
| 1PVH | 129 | 2401 | train | 2530 | 5,1 % |
| 1PXV | 99 | 1377 | train | 1476 | 6,7 % |
| 1QA9 | 258 | 2258 | train | 2516 | 10,3 % |
| 1R0R | 273 | 2257 | train | 2530 | 10,8 % |
| 1R6Q | 146 | 2384 | train | 2530 | 5,8 % |
| 1R8S | 10 | 1543 | train | 1553 | 0,6 % |
| 1RKE | 10 | 1264 | train | 1274 | 0,8 % |
| 1S1Q | 228 | 2302 | train | 2530 | 9,0 % |
| 1SBB | 132 | 941 | train | 1073 | 12,3 % |
| 1SYX | 271 | 2259 | train | 2530 | 10,7 % |
| 1T6B | 240 | 1307 | train | 1547 | 15,5 % |
| 1TMQ | 236 | 2294 | train | 2530 | 9,3 % |
| 1UDI | 33 | 2496 | train | 2529 | 1,3 % |
| 1US7 | 60 | 931 | eval | 991 | 6,1 % |
| 1WQ1 | 14 | 2375 | train | 2389 | 0,6 % |
| 1XD3 | 293 | 2212 | eval | 2505 | 11,7 % |
| 1Y64 | 10 | 1404 | train | 1414 | 0,7 % |
| 1YVB | 159 | 2371 | train | 2530 | 6,3 % |
| 1Z0K | 179 | 2351 | eval | 2530 | 7,1 % |
| 1Z5Y | 123 | 2407 | train | 2530 | 4,9 % |
| 1ZHH | 240 | 2290 | eval | 2530 | 9,5 % |
| 1ZHI | 149 | 2381 | train | 2530 | 5,9 % |
| 1ZLI | 48 | 267 | eval | 315 | 15,2 % |
| 1ZM4 | 233 | 45 | eval | 278 | 83,8 % |
| 2A1A | 172 | 2127 | eval | 2299 | 7,5 % |
| 2A5T | 100 | 1906 | eval | 2006 | 5,0 % |
| 2A9K | 46 | 2484 | eval | 2530 | 1,8 % |
| 2ABZ | 157 | 1839 | train | 1996 | 7,9 % |
| 2AJF | 208 | 493 | eval | 701 | 29,7 % |
| 2AYO | 38 | 2492 | train | 2530 | 1,5 % |
| 2B42 | 16 | 2463 | train | 2479 | 0,6 % |
| 2BTF | 171 | 2317 | train | 2488 | 6,9 % |
| 2C0L | 12 | 1408 | train | 1420 | 0,8 % |
| 2CFH | 156 | 524 | eval | 680 | 22,9 % |
| 2FJU | 293 | 1745 | eval | 2038 | 14,4 % |
| 2G77 | 79 | 2451 | train | 2530 | 3,1 % |
| 2GAF | 33 | 1215 | train | 1248 | 2,6 % |
| 2GTP | 175 | 2355 | train | 2530 | 6,9 % |
| 2H7V | 104 | 2425 | train | 2529 | 4,1 % |
| 2HLE | 105 | 2425 | train | 2530 | 4,2 % |
| 2HQS | 192 | 2338 | train | 2530 | 7,6 % |
| 2HRK | 118 | 2412 | train | 2530 | 4,7 % |
| 2I25 | 123 | 2407 | train | 2530 | 4,9 % |
| 2I9B | 16 | 2514 | eval | 2530 | 0,6 % |
| 2IDO | 30 | 2500 | train | 2530 | 1,2 % |
| 2J0T | 207 | 2323 | train | 2530 | 8,2 % |
| 2J7P | 10 | 1534 | eval | 1544 | 0,6 % |
| 2NZ8 | 10 | 1045 | eval | 1055 | 0,9 % |
| 2O3B | 12 | 1764 | train | 1776 | 0,7 % |
| 2O8V | 216 | 2314 | eval | 2530 | 8,5 % |
| 2OOB | 234 | 2296 | train | 2530 | 9,2 % |
| 2OT3 | 10 | 2520 | eval | 2530 | 0,4 % |
| 2OUL | 295 | 2231 | train | 2526 | 11,7 % |
| 2PCC | 69 | 2461 | train | 2530 | 2,7 % |
| 2SIC | 271 | 2259 | train | 2530 | 10,7 % |
| 2UUY | 197 | 2333 | train | 2530 | 7,8 % |
| 2VDB | 236 | 1423 | train | 1659 | 14,2 % |
| 2X9A | 111 | 2419 | train | 2530 | 4,4 % |
| 3A4S | 80 | 1988 | train | 2068 | 3,9 % |
| 3AAD | 10 | 2225 | train | 2235 | 0,4 % |
| 3BIW | 168 | 2362 | train | 2530 | 6,6 % |
| 3BX7 | 18 | 2512 | train | 2530 | 0,7 % |
| 3CPH | 18 | 2512 | train | 2530 | 0,7 % |
| 3D5S | 201 | 2329 | train | 2530 | 7,9 % |
| 3DAW | 146 | 2278 | eval | 2424 | 6,0 % |
| 3F1P | 94 | 2436 | train | 2530 | 3,7 % |
| 3FN1 | 10 | 2520 | train | 2530 | 0,4 % |
| 3H2V | 136 | 2394 | train | 2530 | 5,4 % |
| 3PC8 | 271 | 2259 | train | 2530 | 10,7 % |
| 3S9D | 31 | 2343 | train | 2374 | 1,3 % |
| 3SGQ | 114 | 2416 | train | 2530 | 4,5 % |
| 3VLB | 269 | 2261 | train | 2530 | 10,6 % |
| 4CPA | 286 | 2244 | train | 2530 | 11,3 % |
| 4FZA | 18 | 2512 | train | 2530 | 0,7 % |
| 4H03 | 201 | 1461 | train | 1662 | 12,1 % |
| 4IZ7 | 10 | 2507 | train | 2517 | 0,4 % |
| 4M76 | 321 | 2209 | train | 2530 | 12,7 % |
| 7CEI | 229 | 2301 | train | 2530 | 9,1 % |
| BAAD | 166 | 2364 | train | 2530 | 6,6 % |
| BOYV | 272 | 2258 | train | 2530 | 10,8 % |

Table S3 Training and evaluation set composition – fold2

| fold2 | fnat ≥ 0.3 | fnat < 0.3 | set | total | % good models |
| --- | --- | --- | --- | --- | --- |
| 1ACB | 20 | 1328 | eval | 1348 | 1,5 % |
| 1ATN | 33 | 2063 | train | 2096 | 1,6 % |
| 1AVX | 163 | 1635 | eval | 1798 | 9,1 % |
| 1AY7 | 246 | 2284 | train | 2530 | 9,7 % |
| 1B6C | 88 | 2442 | train | 2530 | 3,5 % |
| 1BKD | 12 | 1743 | train | 1755 | 0,7 % |
| 1BUH | 208 | 2299 | train | 2507 | 8,3 % |
| 1CLV | 17 | 2513 | train | 2530 | 0,7 % |
| 1D6R | 89 | 2441 | eval | 2530 | 3,5 % |
| 1DFJ | 33 | 2400 | train | 2433 | 1,4 % |
| 1E6E | 286 | 2243 | train | 2529 | 11,3 % |
| 1E96 | 293 | 2237 | train | 2530 | 11,6 % |
| 1EAW | 108 | 2395 | train | 2503 | 4,3 % |
| 1EFN | 138 | 2392 | train | 2530 | 5,5 % |
| 1EWY | 163 | 2367 | train | 2530 | 6,4 % |
| 1F34 | 115 | 2414 | train | 2529 | 4,5 % |
| 1FC2 | 97 | 2433 | eval | 2530 | 3,8 % |
| 1FFW | 209 | 2321 | eval | 2530 | 8,3 % |
| 1FLE | 122 | 2408 | train | 2530 | 4,8 % |
| 1FQ1 | 111 | 2419 | train | 2530 | 4,4 % |
| 1FQJ | 178 | 2352 | train | 2530 | 7,0 % |
| 1GCQ | 269 | 2261 | eval | 2530 | 10,6 % |
| 1GHQ | 127 | 2403 | train | 2530 | 5,0 % |
| 1GL1 | 84 | 2446 | train | 2530 | 3,3 % |
| 1GLA | 22 | 2508 | train | 2530 | 0,9 % |
| 1GPW | 69 | 2461 | train | 2530 | 2,7 % |
| 1GRN | 188 | 516 | train | 704 | 26,7 % |
| 1GXD | 209 | 2321 | train | 2530 | 8,3 % |
| 1H9D | 38 | 2492 | train | 2530 | 1,5 % |
| 1HE1 | 173 | 1969 | train | 2142 | 8,1 % |
| 1HE8 | 106 | 2424 | train | 2530 | 4,2 % |
| 1I2M | 11 | 2519 | train | 2530 | 0,4 % |
| 1J2J | 270 | 2260 | train | 2530 | 10,7 % |
| 1JIW | 271 | 792 | train | 1063 | 25,5 % |
| 1JK9 | 259 | 2271 | train | 2530 | 10,2 % |
| 1JTD | 146 | 2323 | train | 2469 | 5,9 % |
| 1JTG | 107 | 2423 | train | 2530 | 4,2 % |
| 1KAC | 153 | 2377 | train | 2530 | 6,0 % |
| 1KTZ | 122 | 2408 | train | 2530 | 4,8 % |
| 1KXP | 163 | 2367 | eval | 2530 | 6,4 % |
| 1KXQ | 251 | 1666 | train | 1917 | 13,1 % |
| 1LFD | 101 | 2429 | eval | 2530 | 4,0 % |
| 1M10 | 47 | 2483 | train | 2530 | 1,9 % |
| 1MAH | 301 | 2229 | train | 2530 | 11,9 % |
| 1MQ8 | 278 | 2252 | train | 2530 | 11,0 % |
| 1NW9 | 36 | 2494 | train | 2530 | 1,4 % |
| 1OC0 | 81 | 1007 | train | 1088 | 7,4 % |
| 1PVH | 140 | 2390 | train | 2530 | 5,5 % |
| 1PXV | 157 | 1362 | train | 1519 | 10,3 % |
| 1QA9 | 252 | 2278 | train | 2530 | 10,0 % |
| 1R0R | 278 | 2252 | eval | 2530 | 11,0 % |
| 1R6Q | 198 | 2332 | train | 2530 | 7,8 % |
| 1R8S | 10 | 1583 | eval | 1593 | 0,6 % |
| 1RKE | 11 | 1252 | train | 1263 | 0,9 % |
| 1S1Q | 232 | 2298 | train | 2530 | 9,2 % |
| 1SBB | 135 | 1028 | train | 1163 | 11,6 % |
| 1SYX | 280 | 2250 | train | 2530 | 11,1 % |
| 1T6B | 233 | 1314 | train | 1547 | 15,1 % |
| 1TMQ | 238 | 2292 | train | 2530 | 9,4 % |
| 1UDI | 74 | 2456 | train | 2530 | 2,9 % |
| 1US7 | 85 | 917 | train | 1002 | 8,5 % |
| 1WQ1 | 14 | 2350 | train | 2364 | 0,6 % |
| 1XD3 | 328 | 2202 | train | 2530 | 13,0 % |
| 1Y64 | 10 | 1418 | train | 1428 | 0,7 % |
| 1YVB | 166 | 2364 | train | 2530 | 6,6 % |
| 1Z0K | 176 | 2354 | train | 2530 | 7,0 % |
| 1Z5Y | 129 | 2401 | train | 2530 | 5,1 % |
| 1ZHH | 271 | 2259 | train | 2530 | 10,7 % |
| 1ZHI | 135 | 2395 | train | 2530 | 5,3 % |
| 1ZLI | 47 | 334 | train | 381 | 12,3 % |
| 1ZM4 | 239 | 54 | train | 293 | 81,6 % |
| 2A1A | 186 | 2128 | train | 2314 | 8,0 % |
| 2A5T | 110 | 1935 | train | 2045 | 5,4 % |
| 2A9K | 51 | 2479 | train | 2530 | 2,0 % |
| 2ABZ | 163 | 1868 | train | 2031 | 8,0 % |
| 2AJF | 210 | 489 | train | 699 | 30,0 % |
| 2AYO | 38 | 2492 | train | 2530 | 1,5 % |
| 2B42 | 122 | 2353 | train | 2475 | 4,9 % |
| 2BTF | 197 | 2287 | eval | 2484 | 7,9 % |
| 2C0L | 71 | 1338 | eval | 1409 | 5,0 % |
| 2CFH | 156 | 560 | train | 716 | 21,8 % |
| 2FJU | 294 | 1739 | train | 2033 | 14,5 % |
| 2G77 | 73 | 2457 | train | 2530 | 2,9 % |
| 2GAF | 41 | 1175 | train | 1216 | 3,4 % |
| 2GTP | 182 | 2343 | eval | 2525 | 7,2 % |
| 2H7V | 99 | 2431 | train | 2530 | 3,9 % |
| 2HLE | 117 | 2413 | train | 2530 | 4,6 % |
| 2HQS | 180 | 2350 | eval | 2530 | 7,1 % |
| 2HRK | 61 | 2405 | eval | 2466 | 2,5 % |
| 2I25 | 120 | 2410 | train | 2530 | 4,7 % |
| 2I9B | 60 | 2470 | train | 2530 | 2,4 % |
| 2IDO | 59 | 2471 | train | 2530 | 2,3 % |
| 2J0T | 208 | 2322 | eval | 2530 | 8,2 % |
| 2J7P | 13 | 1491 | train | 1504 | 0,9 % |
| 2NZ8 | 20 | 1054 | train | 1074 | 1,9 % |
| 2O3B | 16 | 1779 | train | 1795 | 0,9 % |
| 2O8V | 236 | 2294 | train | 2530 | 9,3 % |
| 2OOB | 251 | 2279 | train | 2530 | 9,9 % |
| 2OT3 | 10 | 2520 | train | 2530 | 0,4 % |
| 2OUL | 297 | 2231 | train | 2528 | 11,7 % |
| 2PCC | 86 | 2444 | eval | 2530 | 3,4 % |
| 2SIC | 278 | 2252 | train | 2530 | 11,0 % |
| 2UUY | 281 | 2249 | eval | 2530 | 11,1 % |
| 2VDB | 228 | 1421 | eval | 1649 | 13,8 % |
| 2X9A | 209 | 2321 | train | 2530 | 8,3 % |
| 3A4S | 75 | 2023 | train | 2098 | 3,6 % |
| 3AAD | 10 | 2190 | train | 2200 | 0,5 % |
| 3BIW | 184 | 2346 | train | 2530 | 7,3 % |
| 3BX7 | 18 | 2509 | train | 2527 | 0,7 % |
| 3CPH | 21 | 2503 | train | 2524 | 0,8 % |
| 3D5S | 253 | 2277 | train | 2530 | 10,0 % |
| 3DAW | 141 | 2299 | train | 2440 | 5,8 % |
| 3F1P | 105 | 2425 | train | 2530 | 4,2 % |
| 3FN1 | 10 | 2520 | eval | 2530 | 0,4 % |
| 3H2V | 144 | 2386 | eval | 2530 | 5,7 % |
| 3PC8 | 284 | 2246 | train | 2530 | 11,2 % |
| 3S9D | 39 | 2325 | eval | 2364 | 1,6 % |
| 3SGQ | 133 | 2397 | train | 2530 | 5,3 % |
| 3VLB | 275 | 2255 | train | 2530 | 10,9 % |
| 4CPA | 284 | 2246 | eval | 2530 | 11,2 % |
| 4FZA | 43 | 2487 | eval | 2530 | 1,7 % |
| 4H03 | 224 | 1448 | eval | 1672 | 13,4 % |
| 4IZ7 | 12 | 2518 | train | 2530 | 0,5 % |
| 4M76 | 344 | 2186 | train | 2530 | 13,6 % |
| 7CEI | 223 | 2307 | train | 2530 | 8,8 % |
| BAAD | 162 | 2368 | train | 2530 | 6,4 % |
| BOYV | 269 | 2261 | train | 2530 | 10,6 % |

Table S4 Training and evaluation set composition – fold3

| fold3 | fnat ≥ 0.3 | fnat < 0.3 | set | total | % good models |
| --- | --- | --- | --- | --- | --- |
| 1ACB | 20 | 1331 | train | 1351 | 1,5 % |
| 1ATN | 39 | 2084 | train | 2123 | 1,8 % |
| 1AVX | 164 | 1591 | train | 1755 | 9,3 % |
| 1AY7 | 244 | 2286 | train | 2530 | 9,6 % |
| 1B6C | 128 | 2402 | train | 2530 | 5,1 % |
| 1BKD | 14 | 1732 | eval | 1746 | 0,8 % |
| 1BUH | 208 | 2317 | eval | 2525 | 8,2 % |
| 1CLV | 60 | 2470 | eval | 2530 | 2,4 % |
| 1D6R | 90 | 2440 | train | 2530 | 3,6 % |
| 1DFJ | 252 | 2200 | train | 2452 | 10,3 % |
| 1E96 | 289 | 2239 | train | 2528 | 11,4 % |
| 1E6E | 294 | 2236 | train | 2530 | 11,6 % |
| 1EAW | 109 | 2397 | eval | 2506 | 4,3 % |
| 1EFN | 179 | 2351 | train | 2530 | 7,1 % |
| 1EWY | 165 | 2365 | train | 2530 | 6,5 % |
| 1F34 | 171 | 2359 | eval | 2530 | 6,8 % |
| 1FC2 | 112 | 2418 | train | 2530 | 4,4 % |
| 1FFW | 213 | 2317 | train | 2530 | 8,4 % |
| 1FLE | 176 | 2354 | train | 2530 | 7,0 % |
| 1FQ1 | 133 | 2397 | train | 2530 | 5,3 % |
| 1FQJ | 244 | 2286 | train | 2530 | 9,6 % |
| 1GCQ | 266 | 2264 | train | 2530 | 10,5 % |
| 1GHQ | 124 | 2406 | train | 2530 | 4,9 % |
| 1GL1 | 83 | 2447 | train | 2530 | 3,3 % |
| 1GLA | 125 | 2405 | train | 2530 | 4,9 % |
| 1GPW | 87 | 2443 | train | 2530 | 3,4 % |
| 1GRN | 237 | 470 | train | 707 | 33,5 % |
| 1GXD | 217 | 2313 | train | 2530 | 8,6 % |
| 1H9D | 119 | 2411 | eval | 2530 | 4,7 % |
| 1HE1 | 59 | 2109 | train | 2168 | 2,7 % |
| 1HE8 | 122 | 2408 | train | 2530 | 4,8 % |
| 1I2M | 10 | 2520 | eval | 2530 | 0,4 % |
| 1J2J | 276 | 2254 | train | 2530 | 10,9 % |
| 1JIW | 283 | 846 | train | 1129 | 25,1 % |
| 1JK9 | 274 | 2256 | train | 2530 | 10,8 % |
| 1JTD | 164 | 2307 | train | 2471 | 6,6 % |
| 1JTG | 108 | 2422 | eval | 2530 | 4,3 % |
| 1KAC | 166 | 2364 | train | 2530 | 6,6 % |
| 1KTZ | 123 | 2407 | train | 2530 | 4,9 % |
| 1KXP | 163 | 2367 | train | 2530 | 6,4 % |
| 1KXQ | 248 | 1718 | train | 1966 | 12,6 % |
| 1LFD | 90 | 2440 | train | 2530 | 3,6 % |
| 1M10 | 114 | 2416 | train | 2530 | 4,5 % |
| 1MAH | 297 | 2233 | train | 2530 | 11,7 % |
| 1MQ8 | 279 | 2251 | train | 2530 | 11,0 % |
| 1NW9 | 78 | 2452 | train | 2530 | 3,1 % |
| 1OC0 | 110 | 951 | eval | 1061 | 10,4 % |
| 1PVH | 136 | 2299 | eval | 2435 | 5,6 % |
| 1PXV | 10 | 1477 | train | 1487 | 0,7 % |
| 1QA9 | 276 | 2254 | train | 2530 | 10,9 % |
| 1R0R | 274 | 2256 | train | 2530 | 10,8 % |
| 1R6Q | 229 | 2301 | eval | 2530 | 9,1 % |
| 1R8S | 10 | 1557 | train | 1567 | 0,6 % |
| 1RKE | 15 | 1249 | eval | 1264 | 1,2 % |
| 1S1Q | 231 | 2299 | eval | 2530 | 9,1 % |
| 1SBB | 134 | 994 | train | 1128 | 11,9 % |
| 1SYX | 284 | 2246 | train | 2530 | 11,2 % |
| 1T6B | 204 | 1307 | train | 1511 | 13,5 % |
| 1TMQ | 297 | 2233 | train | 2530 | 11,7 % |
| 1UDI | 166 | 2364 | eval | 2530 | 6,6 % |
| 1US7 | 59 | 906 | train | 965 | 6,1 % |
| 1WQ1 | 15 | 2366 | eval | 2381 | 0,6 % |
| 1XD3 | 328 | 2202 | train | 2530 | 13,0 % |
| 1Y64 | 10 | 1389 | eval | 1399 | 0,7 % |
| 1YVB | 166 | 2364 | eval | 2530 | 6,6 % |
| 1Z0K | 171 | 2359 | train | 2530 | 6,8 % |
| 1Z5Y | 153 | 2377 | train | 2530 | 6,0 % |
| 1ZHH | 280 | 2250 | train | 2530 | 11,1 % |
| 1ZHI | 154 | 2376 | train | 2530 | 6,1 % |
| 1ZLI | 55 | 318 | train | 373 | 14,7 % |
| 1ZM4 | 237 | 56 | train | 293 | 80,9 % |
| 2HRK | 10 | 2402 | train | 2412 | 0,4 % |
| 2A1A | 215 | 2108 | train | 2323 | 9,3 % |
| 2A5T | 142 | 1936 | train | 2078 | 6,8 % |
| 2A9K | 51 | 2479 | train | 2530 | 2,0 % |
| 2ABZ | 194 | 1823 | eval | 2017 | 9,6 % |
| 2AJF | 223 | 499 | train | 722 | 30,9 % |
| 2AYO | 167 | 2363 | eval | 2530 | 6,6 % |
| 2B42 | 215 | 2268 | eval | 2483 | 8,7 % |
| 2BTF | 225 | 2253 | train | 2478 | 9,1 % |
| 2C0L | 10 | 1432 | train | 1442 | 0,7 % |
| 2CFH | 156 | 536 | train | 692 | 22,5 % |
| 2FJU | 300 | 1728 | train | 2028 | 14,8 % |
| 2G77 | 106 | 2424 | train | 2530 | 4,2 % |
| 2GAF | 82 | 1195 | train | 1277 | 6,4 % |
| 2GTP | 192 | 2338 | train | 2530 | 7,6 % |
| 2H7V | 107 | 2423 | eval | 2530 | 4,2 % |
| 2HLE | 188 | 2342 | train | 2530 | 7,4 % |
| 2HQS | 202 | 2328 | train | 2530 | 8,0 % |
| 2I25 | 174 | 2356 | train | 2530 | 6,9 % |
| 2I9B | 61 | 2469 | train | 2530 | 2,4 % |
| 2IDO | 87 | 2443 | train | 2530 | 3,4 % |
| 2J0T | 277 | 2253 | train | 2530 | 10,9 % |
| 2J7P | 49 | 1453 | train | 1502 | 3,3 % |
| 2NZ8 | 44 | 1006 | train | 1050 | 4,2 % |
| 2O3B | 22 | 1789 | eval | 1811 | 1,2 % |
| 2O8V | 238 | 2292 | train | 2530 | 9,4 % |
| 2OOB | 249 | 2281 | train | 2530 | 9,8 % |
| 2OT3 | 11 | 2519 | train | 2530 | 0,4 % |
| 2OUL | 298 | 2230 | train | 2528 | 11,8 % |
| 2PCC | 106 | 2424 | train | 2530 | 4,2 % |
| 2SIC | 280 | 2250 | train | 2530 | 11,1 % |
| 2UUY | 282 | 2248 | train | 2530 | 11,1 % |
| 2VDB | 254 | 1394 | train | 1648 | 15,4 % |
| 2X9A | 254 | 2276 | train | 2530 | 10,0 % |
| 3A4S | 116 | 1969 | train | 2085 | 5,6 % |
| 3AAD | 10 | 2218 | train | 2228 | 0,4 % |
| 3BIW | 184 | 2346 | train | 2530 | 7,3 % |
| 3BX7 | 149 | 2381 | train | 2530 | 5,9 % |
| 3CPH | 43 | 2487 | eval | 2530 | 1,7 % |
| 3D5S | 250 | 2280 | train | 2530 | 9,9 % |
| 3DAW | 196 | 2239 | train | 2435 | 8,0 % |
| 3F1P | 108 | 2422 | train | 2530 | 4,3 % |
| 3FN1 | 10 | 2520 | train | 2530 | 0,4 % |
| 3H2V | 152 | 2378 | train | 2530 | 6,0 % |
| 3PC8 | 296 | 2234 | train | 2530 | 11,7 % |
| 3S9D | 46 | 2291 | train | 2337 | 2,0 % |
| 3SGQ | 227 | 2303 | train | 2530 | 9,0 % |
| 3VLB | 274 | 2256 | train | 2530 | 10,8 % |
| 4CPA | 275 | 2255 | train | 2530 | 10,9 % |
| 4FZA | 125 | 2405 | train | 2530 | 4,9 % |
| 4H03 | 270 | 1466 | train | 1736 | 15,6 % |
| 4IZ7 | 11 | 2519 | train | 2530 | 0,4 % |
| 4M76 | 332 | 2198 | train | 2530 | 13,1 % |
| 7CEI | 221 | 2309 | eval | 2530 | 8,7 % |
| BAAD | 206 | 2324 | eval | 2530 | 8,1 % |
| BOYV | 269 | 2261 | train | 2530 | 10,6 % |

Table S5 Training and evaluation set composition – fold4

| fold4 | fnat ≥ 0.3 | fnat < 0.3 | set | total | % good models |
| --- | --- | --- | --- | --- | --- |
| 1ACB | 16 | 1309 | train | 1325 | 1,2 % |
| 1ATN | 68 | 2040 | eval | 2108 | 3,2 % |
| 1AVX | 177 | 1574 | train | 1751 | 10,1 % |
| 1AY7 | 247 | 2283 | train | 2530 | 9,8 % |
| 1B6C | 60 | 2369 | eval | 2429 | 2,5 % |
| 1BKD | 15 | 1796 | train | 1811 | 0,8 % |
| 1BUH | 219 | 2311 | train | 2530 | 8,7 % |
| 1CLV | 86 | 2444 | train | 2530 | 3,4 % |
| 1D6R | 98 | 2432 | train | 2530 | 3,9 % |
| 1DFJ | 249 | 2182 | eval | 2431 | 10,2 % |
| 1E6E | 307 | 2222 | train | 2529 | 12,1 % |
| 1E96 | 293 | 2224 | train | 2517 | 11,6 % |
| 1EAW | 107 | 2423 | train | 2530 | 4,2 % |
| 1EFN | 189 | 2341 | train | 2530 | 7,5 % |
| 1EWY | 171 | 2359 | train | 2530 | 6,8 % |
| 1F34 | 307 | 2223 | train | 2530 | 12,1 % |
| 1FC2 | 149 | 2381 | train | 2530 | 5,9 % |
| 1FFW | 211 | 2319 | train | 2530 | 8,3 % |
| 1FLE | 177 | 2353 | eval | 2530 | 7,0 % |
| 1FQ1 | 138 | 2392 | train | 2530 | 5,5 % |
| 1FQJ | 241 | 2289 | train | 2530 | 9,5 % |
| 1GCQ | 267 | 2263 | train | 2530 | 10,6 % |
| 1GHQ | 126 | 2404 | eval | 2530 | 5,0 % |
| 1GL1 | 82 | 2448 | train | 2530 | 3,2 % |
| 1GLA | 288 | 2242 | train | 2530 | 11,4 % |
| 1GPW | 85 | 2445 | train | 2530 | 3,4 % |
| 1GRN | 277 | 458 | train | 735 | 37,7 % |
| 1GXD | 229 | 2301 | eval | 2530 | 9,1 % |
| 1H9D | 130 | 2400 | train | 2530 | 5,1 % |
| 1HE1 | 23 | 2124 | train | 2147 | 1,1 % |
| 1HE8 | 142 | 2388 | train | 2530 | 5,6 % |
| 1I2M | 57 | 2473 | train | 2530 | 2,3 % |
| 1J2J | 296 | 2234 | eval | 2530 | 11,7 % |
| 1JIW | 182 | 834 | train | 1016 | 17,9 % |
| 1JK9 | 277 | 2253 | train | 2530 | 10,9 % |
| 1JTD | 164 | 2310 | train | 2474 | 6,6 % |
| 1JTG | 126 | 2404 | train | 2530 | 5,0 % |
| 1KAC | 218 | 2312 | eval | 2530 | 8,6 % |
| 1KTZ | 124 | 2406 | train | 2530 | 4,9 % |
| 1KXP | 258 | 2249 | train | 2507 | 10,3 % |
| 1KXQ | 252 | 1704 | eval | 1956 | 12,9 % |
| 1LFD | 120 | 2410 | train | 2530 | 4,7 % |
| 1M10 | 21 | 2509 | eval | 2530 | 0,8 % |
| 1MAH | 299 | 2231 | train | 2530 | 11,8 % |
| 1MQ8 | 280 | 2250 | train | 2530 | 11,1 % |
| 1NW9 | 84 | 2446 | train | 2530 | 3,3 % |
| 1OC0 | 123 | 985 | train | 1108 | 11,1 % |
| 1PVH | 87 | 2281 | train | 2368 | 3,7 % |
| 1PXV | 10 | 1532 | eval | 1542 | 0,6 % |
| 1QA9 | 293 | 2237 | train | 2530 | 11,6 % |
| 1R0R | 276 | 2254 | train | 2530 | 10,9 % |
| 1R6Q | 225 | 2305 | train | 2530 | 8,9 % |
| 1R8S | 10 | 1553 | train | 1563 | 0,6 % |
| 1RKE | 18 | 1256 | train | 1274 | 1,4 % |
| 1S1Q | 232 | 2298 | train | 2530 | 9,2 % |
| 1SBB | 156 | 992 | train | 1148 | 13,6 % |
| 1SYX | 299 | 2231 | eval | 2530 | 11,8 % |
| 1T6B | 179 | 1293 | eval | 1472 | 12,2 % |
| 1TMQ | 298 | 2232 | eval | 2530 | 11,8 % |
| 1UDI | 166 | 2355 | train | 2521 | 6,6 % |
| 1US7 | 68 | 934 | train | 1002 | 6,8 % |
| 1WQ1 | 35 | 2341 | train | 2376 | 1,5 % |
| 1XD3 | 340 | 2179 | train | 2519 | 13,5 % |
| 1Y64 | 10 | 1361 | train | 1371 | 0,7 % |
| 1YVB | 166 | 2364 | train | 2530 | 6,6 % |
| 1Z0K | 210 | 2320 | train | 2530 | 8,3 % |
| 1Z5Y | 196 | 2334 | train | 2530 | 7,7 % |
| 1ZHH | 276 | 2254 | train | 2530 | 10,9 % |
| 1ZHI | 187 | 2343 | train | 2530 | 7,4 % |
| 1ZLI | 48 | 331 | train | 379 | 12,7 % |
| 1ZM4 | 238 | 55 | train | 293 | 81,2 % |
| 2A1A | 258 | 2051 | train | 2309 | 11,2 % |
| 2A5T | 144 | 1891 | train | 2035 | 7,1 % |
| 2A9K | 55 | 2475 | train | 2530 | 2,2 % |
| 2ABZ | 264 | 1744 | train | 2008 | 13,1 % |
| 2AJF | 247 | 480 | train | 727 | 34,0 % |
| 2AYO | 179 | 2351 | train | 2530 | 7,1 % |
| 2B42 | 210 | 2265 | train | 2475 | 8,5 % |
| 2BTF | 283 | 2196 | train | 2479 | 11,4 % |
| 2C0L | 11 | 1453 | train | 1464 | 0,8 % |
| 2CFH | 203 | 493 | train | 696 | 29,2 % |
| 2FJU | 303 | 1716 | train | 2019 | 15,0 % |
| 2G77 | 153 | 2377 | eval | 2530 | 6,0 % |
| 2GAF | 100 | 1165 | eval | 1265 | 7,9 % |
| 2GTP | 193 | 2329 | train | 2522 | 7,7 % |
| 2H7V | 131 | 2399 | train | 2530 | 5,2 % |
| 2HLE | 188 | 2342 | train | 2530 | 7,4 % |
| 2HQS | 257 | 2273 | train | 2530 | 10,2 % |
| 2HRK | 10 | 2402 | train | 2412 | 0,4 % |
| 2I25 | 227 | 2303 | train | 2530 | 9,0 % |
| 2I9B | 60 | 2470 | train | 2530 | 2,4 % |
| 2IDO | 120 | 2410 | train | 2530 | 4,7 % |
| 2J0T | 299 | 2231 | train | 2530 | 11,8 % |
| 2J7P | 51 | 1518 | train | 1569 | 3,3 % |
| 2NZ8 | 33 | 1015 | train | 1048 | 3,1 % |
| 2O3B | 20 | 1793 | train | 1813 | 1,1 % |
| 2O8V | 228 | 2302 | train | 2530 | 9,0 % |
| 2OOB | 255 | 2275 | eval | 2530 | 10,1 % |
| 2OT3 | 13 | 2517 | train | 2530 | 0,5 % |
| 2OUL | 298 | 2229 | train | 2527 | 11,8 % |
| 2PCC | 117 | 2413 | train | 2530 | 4,6 % |
| 2SIC | 279 | 2251 | eval | 2530 | 11,0 % |
| 2UUY | 283 | 2247 | train | 2530 | 11,2 % |
| 2VDB | 271 | 1362 | train | 1633 | 16,6 % |
| 2X9A | 103 | 2427 | train | 2530 | 4,1 % |
| 3A4S | 120 | 1939 | train | 2059 | 5,8 % |
| 3AAD | 11 | 2167 | train | 2178 | 0,5 % |
| 3BIW | 175 | 2355 | eval | 2530 | 6,9 % |
| 3BX7 | 184 | 2346 | train | 2530 | 7,3 % |
| 3CPH | 76 | 2454 | train | 2530 | 3,0 % |
| 3D5S | 268 | 2262 | train | 2530 | 10,6 % |
| 3DAW | 225 | 2210 | train | 2435 | 9,2 % |
| 3F1P | 113 | 2417 | eval | 2530 | 4,5 % |
| 3FN1 | 12 | 2518 | train | 2530 | 0,5 % |
| 3H2V | 151 | 2379 | train | 2530 | 6,0 % |
| 3PC8 | 296 | 2234 | eval | 2530 | 11,7 % |
| 3S9D | 55 | 2313 | train | 2368 | 2,3 % |
| 3SGQ | 220 | 2310 | eval | 2530 | 8,7 % |
| 3VLB | 272 | 2258 | eval | 2530 | 10,8 % |
| 4CPA | 274 | 2256 | train | 2530 | 10,8 % |
| 4FZA | 126 | 2404 | train | 2530 | 5,0 % |
| 4H03 | 275 | 1466 | train | 1741 | 15,8 % |
| 4IZ7 | 13 | 2517 | train | 2530 | 0,5 % |
| 4M76 | 334 | 2196 | eval | 2530 | 13,2 % |
| 7CEI | 243 | 2287 | train | 2530 | 9,6 % |
| BAAD | 213 | 2317 | train | 2530 | 8,4 % |
| BOYV | 284 | 2246 | eval | 2530 | 11,2 % |

Table S6 Training and evaluation set composition – fold5

| fold5 | fnat ≥ 0.3 | fnat < 0.3 | set | total | % good models |
| --- | --- | --- | --- | --- | --- |
| 1ACB | 7 | 1803 | train | 1810 | 0,4 % |
| 1ATN | 62 | 2279 | train | 2341 | 2,6 % |
| 1AVX | 212 | 1561 | train | 1773 | 12,0 % |
| 1AY7 | 249 | 2281 | eval | 2530 | 9,8 % |
| 1B6C | 10 | 2317 | train | 2327 | 0,4 % |
| 1BKD | 13 | 2196 | train | 2209 | 0,6 % |
| 1BUH | 227 | 1715 | train | 1942 | 11,7 % |
| 1CLV | 52 | 2478 | train | 2530 | 2,1 % |
| 1D6R | 106 | 2424 | train | 2530 | 4,2 % |
| 1DFJ | 238 | 1832 | train | 2070 | 11,5 % |
| 1E6E | 319 | 2142 | eval | 2461 | 13,0 % |
| 1E96 | 299 | 2051 | eval | 2350 | 12,7 % |
| 1EAW | 110 | 1616 | train | 1726 | 6,4 % |
| 1EFN | 188 | 2342 | eval | 2530 | 7,4 % |
| 1EWY | 167 | 2363 | eval | 2530 | 6,6 % |
| 1F34 | 273 | 2086 | train | 2359 | 11,6 % |
| 1FC2 | 143 | 1984 | train | 2127 | 6,7 % |
| 1FFW | 242 | 2288 | train | 2530 | 9,6 % |
| 1FLE | 147 | 2383 | train | 2530 | 5,8 % |
| 1FQ1 | 137 | 2393 | eval | 2530 | 5,4 % |
| 1FQJ | 244 | 2286 | eval | 2530 | 9,6 % |
| 1GCQ | 264 | 2266 | train | 2530 | 10,4 % |
| 1GHQ | 139 | 2391 | train | 2530 | 5,5 % |
| 1GL1 | 88 | 2442 | eval | 2530 | 3,5 % |
| 1GLA | 278 | 2252 | train | 2530 | 11,0 % |
| 1GPW | 139 | 2391 | eval | 2530 | 5,5 % |
| 1GRN | 270 | 620 | train | 890 | 30,3 % |
| 1GXD | 232 | 2298 | train | 2530 | 9,2 % |
| 1H9D | 141 | 2195 | train | 2336 | 6,0 % |
| 1HE1 | 37 | 2088 | train | 2125 | 1,7 % |
| 1HE8 | 143 | 2387 | train | 2530 | 5,7 % |
| 1I2M | 54 | 2476 | train | 2530 | 2,1 % |
| 1J2J | 316 | 2214 | train | 2530 | 12,5 % |
| 1JIW | 22 | 1079 | train | 1101 | 2,0 % |
| 1JK9 | 274 | 2256 | train | 2530 | 10,8 % |
| 1JTD | 166 | 2172 | eval | 2338 | 7,1 % |
| 1JTG | 125 | 2343 | train | 2468 | 5,1 % |
| 1KAC | 230 | 2300 | train | 2530 | 9,1 % |
| 1KTZ | 128 | 2402 | eval | 2530 | 5,1 % |
| 1KXP | 153 | 2249 | train | 2402 | 6,4 % |
| 1KXQ | 248 | 1809 | train | 2057 | 12,1 % |
| 1LFD | 127 | 2403 | train | 2530 | 5,0 % |
| 1M10 | 12 | 2518 | train | 2530 | 0,5 % |
| 1MAH | 300 | 2230 | eval | 2530 | 11,9 % |
| 1MQ8 | 280 | 2250 | train | 2530 | 11,1 % |
| 1NW9 | 99 | 2431 | train | 2530 | 3,9 % |
| 1OC0 | 55 | 1315 | train | 1370 | 4,0 % |
| 1PVH | 10 | 2271 | train | 2281 | 0,4 % |
| 1PXV | 10 | 1776 | train | 1786 | 0,6 % |
| 1QA9 | 297 | 2144 | eval | 2441 | 12,2 % |
| 1R0R | 279 | 2251 | train | 2530 | 11,0 % |
| 1R6Q | 252 | 2278 | train | 2530 | 10,0 % |
| 1R8S | 10 | 1560 | train | 1570 | 0,6 % |
| 1RKE | 23 | 1271 | train | 1294 | 1,8 % |
| 1S1Q | 232 | 2296 | train | 2528 | 9,2 % |
| 1SBB | 164 | 955 | eval | 1119 | 14,7 % |
| 1SYX | 299 | 2229 | train | 2528 | 11,8 % |
| 1T6B | 54 | 1267 | train | 1321 | 4,1 % |
| 1TMQ | 298 | 2232 | train | 2530 | 11,8 % |
| 1UDI | 94 | 1495 | train | 1589 | 5,9 % |
| 1US7 | 92 | 945 | train | 1037 | 8,9 % |
| 1WQ1 | 37 | 2443 | train | 2480 | 1,5 % |
| 1XD3 | 286 | 2067 | train | 2353 | 12,2 % |
| 1Y64 | 10 | 1408 | train | 1418 | 0,7 % |
| 1YVB | 166 | 2364 | train | 2530 | 6,6 % |
| 1Z0K | 204 | 2326 | train | 2530 | 8,1 % |
| 1Z5Y | 220 | 2310 | eval | 2530 | 8,7 % |
| 1ZHH | 269 | 2261 | train | 2530 | 10,6 % |
| 1ZLI | 48 | 808 | train | 856 | 5,6 % |
| 1ZM4 | 249 | 217 | train | 466 | 53,4 % |
| 2A1A | 261 | 2101 | train | 2362 | 11,0 % |
| 2A5T | 47 | 2177 | train | 2224 | 2,1 % |
| 2A9K | 63 | 2467 | train | 2530 | 2,5 % |
| 2ABZ | 213 | 1794 | train | 2007 | 10,6 % |
| 2AJF | 237 | 918 | train | 1155 | 20,5 % |
| 2B42 | 13 | 2489 | train | 2502 | 0,5 % |
| 2BTF | 261 | 2197 | train | 2458 | 10,6 % |
| 2C0L | 18 | 1386 | train | 1404 | 1,3 % |
| 2CFH | 203 | 1081 | train | 1284 | 15,8 % |
| 2FJU | 301 | 1688 | train | 1989 | 15,1 % |
| 2G77 | 154 | 2376 | train | 2530 | 6,1 % |
| 2GAF | 89 | 1524 | train | 1613 | 5,5 % |
| 2GTP | 197 | 1840 | train | 2037 | 9,7 % |
| 2H7V | 125 | 2330 | train | 2455 | 5,1 % |
| 2HLE | 116 | 2414 | train | 2530 | 4,6 % |
| 2HQS | 260 | 2270 | train | 2530 | 10,3 % |
| 2HRK | 10 | 2402 | train | 2412 | 0,4 % |
| 2I25 | 264 | 2266 | eval | 2530 | 10,4 % |
| 2I9B | 55 | 2475 | train | 2530 | 2,2 % |
| 2IDO | 120 | 2410 | eval | 2530 | 4,7 % |
| 2J0T | 296 | 2146 | train | 2442 | 12,1 % |
| 2J7P | 10 | 2036 | train | 2046 | 0,5 % |
| 2NZ8 | 24 | 1110 | train | 1134 | 2,1 % |
| 2O3B | 30 | 2251 | train | 2281 | 1,3 % |
| 2O8V | 229 | 2301 | train | 2530 | 9,1 % |
| 2OOB | 253 | 2275 | train | 2528 | 10,0 % |
| 2OT3 | 17 | 2513 | train | 2530 | 0,7 % |
| 2OUL | 298 | 2112 | eval | 2410 | 12,4 % |
| 2PCC | 125 | 2405 | train | 2530 | 4,9 % |
| 2SIC | 279 | 2251 | train | 2530 | 11,0 % |
| 2UUY | 316 | 2214 | train | 2530 | 12,5 % |
| 2VDB | 263 | 1361 | train | 1624 | 16,2 % |
| 2X9A | 88 | 2442 | eval | 2530 | 3,5 % |
| 3A4S | 119 | 2081 | eval | 2200 | 5,4 % |
| 3AAD | 15 | 2419 | eval | 2434 | 0,6 % |
| 3BIW | 49 | 2089 | train | 2138 | 2,3 % |
| 3BX7 | 156 | 2183 | eval | 2339 | 6,7 % |
| 3CPH | 96 | 2250 | train | 2346 | 4,1 % |
| 3D5S | 270 | 2260 | eval | 2530 | 10,7 % |
| 3DAW | 229 | 2116 | train | 2345 | 9,8 % |
| 3F1P | 119 | 2344 | train | 2463 | 4,8 % |
| 3FN1 | 12 | 2518 | train | 2530 | 0,5 % |
| 3H2V | 151 | 2379 | train | 2530 | 6,0 % |
| 3PC8 | 305 | 2225 | train | 2530 | 12,1 % |
| 3S9D | 62 | 2381 | train | 2443 | 2,5 % |
| 3SGQ | 91 | 2439 | train | 2530 | 3,6 % |
| 3VLB | 276 | 2254 | train | 2530 | 10,9 % |
| 4CPA | 283 | 2247 | train | 2530 | 11,2 % |
| 4FZA | 38 | 2492 | train | 2530 | 1,5 % |
| 4H03 | 249 | 1557 | train | 1806 | 13,8 % |
| 4IZ7 | 11 | 1753 | eval | 1764 | 0,6 % |
| 4M76 | 334 | 2196 | train | 2530 | 13,2 % |
| 7CEI | 270 | 2260 | train | 2530 | 10,7 % |
| BAAD | 193 | 2337 | train | 2530 | 7,6 % |
| BOYV | 282 | 2248 | train | 2530 | 11,1 % |

Table S7 Training and evaluation set composition – fold6

| fold6 | fnat ≥ 0.3 | fnat < 0.3 | set | total | % good models |
| --- | --- | --- | --- | --- | --- |
| 1ACB | 6 | 1950 | train | 1956 | 0,3 % |
| 1ATN | 58 | 2472 | train | 2530 | 2,3 % |
| 1AVX | 200 | 1477 | train | 1677 | 11,9 % |
| 1AY7 | 252 | 2278 | eval | 2530 | 10,0 % |
| 1B6C | 10 | 2316 | train | 2326 | 0,4 % |
| 1BKD | 13 | 2517 | train | 2530 | 0,5 % |
| 1BUH | 227 | 145 | train | 372 | 61,0 % |
| 1CLV | 13 | 2517 | eval | 2530 | 0,5 % |
| 1D6R | 103 | 2427 | train | 2530 | 4,1 % |
| 1DFJ | 23 | 885 | train | 908 | 2,5 % |
| 1E96 | 304 | 1194 | train | 1498 | 20,3 % |
| 1E6E | 332 | 1918 | eval | 2250 | 14,8 % |
| 1EAW | 117 | 485 | train | 602 | 19,4 % |
| 1EFN | 138 | 2392 | train | 2530 | 5,5 % |
| 1EWY | 184 | 2346 | train | 2530 | 7,3 % |
| 1F34 | 82 | 679 | train | 761 | 10,8 % |
| 1FC2 | 113 | 1015 | train | 1128 | 10,0 % |
| 1FFW | 239 | 2291 | train | 2530 | 9,4 % |
| 1FLE | 155 | 2375 | train | 2530 | 6,1 % |
| 1FQ1 | 126 | 2404 | train | 2530 | 5,0 % |
| 1FQJ | 229 | 2301 | train | 2530 | 9,1 % |
| 1GCQ | 265 | 2265 | train | 2530 | 10,5 % |
| 1GHQ | 149 | 2381 | eval | 2530 | 5,9 % |
| 1GL1 | 98 | 2432 | train | 2530 | 3,9 % |
| 1GLA | 288 | 2242 | train | 2530 | 11,4 % |
| 1GPW | 138 | 2392 | train | 2530 | 5,5 % |
| 1GRN | 162 | 1533 | train | 1695 | 9,6 % |
| 1GXD | 239 | 2291 | eval | 2530 | 9,4 % |
| 1H9D | 148 | 1789 | train | 1937 | 7,6 % |
| 1HE1 | 63 | 2099 | train | 2162 | 2,9 % |
| 1HE8 | 142 | 2388 | eval | 2530 | 5,6 % |
| 1I2M | 10 | 2520 | eval | 2530 | 0,4 % |
| 1J2J | 309 | 2221 | train | 2530 | 12,2 % |
| 1JIW | 22 | 2243 | eval | 2265 | 1,0 % |
| 1JK9 | 277 | 2253 | train | 2530 | 10,9 % |
| 1JTD | 164 | 2366 | train | 2530 | 6,5 % |
| 1JTG | 125 | 2405 | train | 2530 | 4,9 % |
| 1KAC | 224 | 2306 | eval | 2530 | 8,9 % |
| 1KTZ | 127 | 2403 | train | 2530 | 5,0 % |
| 1KXP | 40 | 2249 | train | 2289 | 1,7 % |
| 1KXQ | 247 | 2283 | eval | 2530 | 9,8 % |
| 1LFD | 117 | 2413 | eval | 2530 | 4,6 % |
| 1M10 | 13 | 2517 | eval | 2530 | 0,5 % |
| 1MAH | 309 | 2221 | train | 2530 | 12,2 % |
| 1MQ8 | 280 | 2250 | train | 2530 | 11,1 % |
| 1NW9 | 103 | 2427 | train | 2530 | 4,1 % |
| 1OC0 | 54 | 1688 | train | 1742 | 3,1 % |
| 1PVH | 10 | 2271 | train | 2281 | 0,4 % |
| 1PXV | 117 | 2210 | train | 2327 | 5,0 % |
| 1QA9 | 289 | 1247 | eval | 1536 | 18,8 % |
| 1R0R | 277 | 2253 | train | 2530 | 10,9 % |
| 1R6Q | 253 | 2277 | train | 2530 | 10,0 % |
| 1R8S | 10 | 1578 | eval | 1588 | 0,6 % |
| 1RKE | 10 | 1378 | train | 1388 | 0,7 % |
| 1S1Q | 234 | 2296 | train | 2530 | 9,2 % |
| 1SBB | 169 | 960 | train | 1129 | 15,0 % |
| 1SYX | 258 | 2272 | train | 2530 | 10,2 % |
| 1T6B | 33 | 1307 | train | 1340 | 2,5 % |
| 1TMQ | 307 | 2223 | train | 2530 | 12,1 % |
| 1UDI | 36 | 343 | train | 379 | 9,5 % |
| 1US7 | 87 | 926 | train | 1013 | 8,6 % |
| 1WQ1 | 30 | 2500 | train | 2530 | 1,2 % |
| 1XD3 | 10 | 73 | eval | 83 | 12,0 % |
| 1Y64 | 10 | 1401 | eval | 1411 | 0,7 % |
| 1YVB | 166 | 2364 | train | 2530 | 6,6 % |
| 1Z0K | 212 | 2318 | train | 2530 | 8,4 % |
| 1Z5Y | 191 | 2339 | train | 2530 | 7,5 % |
| 1ZHH | 262 | 2268 | train | 2530 | 10,4 % |
| 1ZHI | 169 | 2361 | train | 2530 | 6,7 % |
| 1ZLI | 48 | 1425 | train | 1473 | 3,3 % |
| 1ZM4 | 258 | 1100 | train | 1358 | 19,0 % |
| 2A1A | 260 | 2270 | train | 2530 | 10,3 % |
| 2A5T | 29 | 2501 | train | 2530 | 1,1 % |
| 2A9K | 71 | 2459 | train | 2530 | 2,8 % |
| 2ABZ | 153 | 1864 | train | 2017 | 7,6 % |
| 2AJF | 240 | 2290 | train | 2530 | 9,5 % |
| 2AYO | 35 | 2495 | train | 2530 | 1,4 % |
| 2B42 | 12 | 2518 | eval | 2530 | 0,5 % |
| 2BTF | 230 | 2102 | train | 2332 | 9,9 % |
| 2C0L | 31 | 1472 | train | 1503 | 2,1 % |
| 2CFH | 204 | 2326 | train | 2530 | 8,1 % |
| 2FJU | 309 | 2220 | train | 2529 | 12,2 % |
| 2G77 | 114 | 2416 | train | 2530 | 4,5 % |
| 2GAF | 141 | 2345 | train | 2486 | 5,7 % |
| 2GTP | 192 | 958 | train | 1150 | 16,7 % |
| 2H7V | 52 | 2222 | train | 2274 | 2,3 % |
| 2HLE | 98 | 2432 | train | 2530 | 3,9 % |
| 2HQS | 229 | 2301 | train | 2530 | 9,1 % |
| 2HRK | 10 | 2402 | train | 2412 | 0,4 % |
| 2I25 | 288 | 2242 | train | 2530 | 11,4 % |
| 2I9B | 82 | 2448 | train | 2530 | 3,2 % |
| 2IDO | 89 | 2441 | train | 2530 | 3,5 % |
| 2J0T | 296 | 1755 | train | 2051 | 14,4 % |
| 2J7P | 11 | 2519 | train | 2530 | 0,4 % |
| 2NZ8 | 26 | 1174 | eval | 1200 | 2,2 % |
| 2O3B | 36 | 2494 | train | 2530 | 1,4 % |
| 2O8V | 229 | 2301 | train | 2530 | 9,1 % |
| 2OOB | 297 | 2233 | train | 2530 | 11,7 % |
| 2OT3 | 17 | 2513 | train | 2530 | 0,7 % |
| 2OUL | 300 | 1329 | train | 1629 | 18,4 % |
| 2PCC | 129 | 2401 | eval | 2530 | 5,1 % |
| 2SIC | 278 | 2252 | train | 2530 | 11,0 % |
| 2UUY | 316 | 2214 | eval | 2530 | 12,5 % |
| 2VDB | 263 | 1384 | train | 1647 | 16,0 % |
| 2X9A | 81 | 2449 | train | 2530 | 3,2 % |
| 3A4S | 133 | 2338 | train | 2471 | 5,4 % |
| 3AAD | 17 | 2513 | train | 2530 | 0,7 % |
| 3BIW | 37 | 2350 | eval | 2387 | 1,6 % |
| 3BX7 | 35 | 1979 | eval | 2014 | 1,7 % |
| 3CPH | 35 | 2035 | train | 2070 | 1,7 % |
| 3D5S | 262 | 2268 | train | 2530 | 10,4 % |
| 3DAW | 163 | 1995 | train | 2158 | 7,6 % |
| 3F1P | 101 | 2281 | train | 2382 | 4,2 % |
| 3FN1 | 22 | 2508 | train | 2530 | 0,9 % |
| 3H2V | 150 | 2380 | train | 2530 | 5,9 % |
| 3PC8 | 328 | 2202 | train | 2530 | 13,0 % |
| 3S9D | 52 | 2478 | eval | 2530 | 2,1 % |
| 3SGQ | 91 | 2439 | train | 2530 | 3,6 % |
| 3VLB | 277 | 2253 | train | 2530 | 10,9 % |
| 4CPA | 292 | 2238 | train | 2530 | 11,5 % |
| 4FZA | 33 | 2497 | train | 2530 | 1,3 % |
| 4H03 | 275 | 2255 | train | 2530 | 10,9 % |
| 4IZ7 | 13 | 1089 | train | 1102 | 1,2 % |
| 4M76 | 337 | 2193 | eval | 2530 | 13,3 % |
| 7CEI | 237 | 2293 | train | 2530 | 9,4 % |
| BAAD | 186 | 2344 | eval | 2530 | 7,4 % |
| BOYV | 282 | 2248 | train | 2530 | 11,1 % |
| eval | dat | a for fold6 |  |  |  |

Table S8 Training and evaluation set composition – fold7

| fold7 | fnat ≥ 0.3 | fnat < 0.3 | set | total | % good models |
| --- | --- | --- | --- | --- | --- |
| 1ACB | 35 | 1938 | eval | 1973 | 1,8 % |
| 1ATN | 83 | 2447 | train | 2530 | 3,3 % |
| 1AVX | 161 | 1520 | train | 1681 | 9,6 % |
| 1AY7 | 268 | 2262 | train | 2530 | 10,6 % |
| 1B6C | 10 | 2316 | eval | 2326 | 0,4 % |
| 1BKD | 17 | 2513 | train | 2530 | 0,7 % |
| 1BUH | 225 | 132 | train | 357 | 63,0 % |
| 1CLV | 13 | 2517 | train | 2530 | 0,5 % |
| 1D6R | 106 | 2424 | train | 2530 | 4,2 % |
| 1DFJ | 23 | 935 | eval | 958 | 2,4 % |
| 1E6E | 322 | 1907 | train | 2229 | 14,4 % |
| 1E96 | 308 | 1150 | train | 1458 | 21,1 % |
| 1EAW | 123 | 430 | train | 553 | 22,2 % |
| 1EFN | 106 | 2424 | train | 2530 | 4,2 % |
| 1EWY | 197 | 2333 | eval | 2530 | 7,8 % |
| 1F34 | 146 | 648 | train | 794 | 18,4 % |
| 1FC2 | 101 | 1058 | train | 1159 | 8,7 % |
| 1FFW | 250 | 2280 | train | 2530 | 9,9 % |
| 1FLE | 156 | 2374 | train | 2530 | 6,2 % |
| 1FQ1 | 129 | 2401 | train | 2530 | 5,1 % |
| 1FQJ | 241 | 2289 | train | 2530 | 9,5 % |
| 1GCQ | 268 | 2262 | train | 2530 | 10,6 % |
| 1GHQ | 143 | 2387 | train | 2530 | 5,7 % |
| 1GL1 | 114 | 2416 | train | 2530 | 4,5 % |
| 1GLA | 161 | 2369 | train | 2530 | 6,4 % |
| 1GPW | 142 | 2388 | train | 2530 | 5,6 % |
| 1GRN | 162 | 1558 | eval | 1720 | 9,4 % |
| 1GXD | 242 | 2288 | train | 2530 | 9,6 % |
| 1H9D | 138 | 1818 | train | 1956 | 7,1 % |
| 1HE1 | 210 | 1904 | train | 2114 | 9,9 % |
| 1HE8 | 142 | 2388 | train | 2530 | 5,6 % |
| 1I2M | 12 | 2518 | train | 2530 | 0,5 % |
| 1J2J | 304 | 2226 | train | 2530 | 12,0 % |
| 1JIW | 25 | 2239 | train | 2264 | 1,1 % |
| 1JK9 | 275 | 2255 | train | 2530 | 10,9 % |
| 1JTD | 165 | 2365 | train | 2530 | 6,5 % |
| 1JTG | 122 | 2408 | train | 2530 | 4,8 % |
| 1KAC | 235 | 2295 | train | 2530 | 9,3 % |
| 1KTZ | 129 | 2401 | train | 2530 | 5,1 % |
| 1KXP | 14 | 2234 | train | 2248 | 0,6 % |
| 1KXQ | 262 | 2268 | train | 2530 | 10,4 % |
| 1LFD | 160 | 2370 | train | 2530 | 6,3 % |
| 1M10 | 61 | 2469 | train | 2530 | 2,4 % |
| 1MAH | 315 | 2215 | train | 2530 | 12,5 % |
| 1MQ8 | 279 | 2251 | eval | 2530 | 11,0 % |
| 1NW9 | 23 | 2507 | train | 2530 | 0,9 % |
| 1OC0 | 94 | 1587 | train | 1681 | 5,6 % |
| 1PVH | 10 | 2271 | train | 2281 | 0,4 % |
| 1PXV | 199 | 2126 | eval | 2325 | 8,6 % |
| 1QA9 | 307 | 1231 | train | 1538 | 20,0 % |
| 1R0R | 277 | 2253 | eval | 2530 | 10,9 % |
| 1R6Q | 241 | 2289 | train | 2530 | 9,5 % |
| 1R8S | 10 | 1608 | train | 1618 | 0,6 % |
| 1RKE | 14 | 1350 | train | 1364 | 1,0 % |
| 1S1Q | 233 | 2297 | train | 2530 | 9,2 % |
| 1SBB | 162 | 1013 | eval | 1175 | 13,8 % |
| 1SYX | 253 | 2277 | eval | 2530 | 10,0 % |
| 1T6B | 33 | 1280 | train | 1313 | 2,5 % |
| 1TMQ | 298 | 2232 | train | 2530 | 11,8 % |
| 1UDI | 36 | 323 | train | 359 | 10,0 % |
| 1US7 | 80 | 883 | eval | 963 | 8,3 % |
| 1WQ1 | 27 | 2503 | train | 2530 | 1,1 % |
| 1XD3 | 13 | 98 | train | 111 | 11,7 % |
| 1Y64 | 10 | 1392 | train | 1402 | 0,7 % |
| 1YVB | 168 | 2362 | train | 2530 | 6,6 % |
| 1Z0K | 213 | 2317 | train | 2530 | 8,4 % |
| 1Z5Y | 159 | 2371 | eval | 2530 | 6,3 % |
| 1ZHH | 272 | 2258 | eval | 2530 | 10,8 % |
| 1ZHI | 161 | 2369 | train | 2530 | 6,4 % |
| 1ZLI | 49 | 1380 | train | 1429 | 3,4 % |
| 1ZM4 | 271 | 1125 | train | 1396 | 19,4 % |
| 2A1A | 198 | 2332 | eval | 2530 | 7,8 % |
| 2A5T | 30 | 2500 | eval | 2530 | 1,2 % |
| 2A9K | 53 | 2477 | eval | 2530 | 2,1 % |
| 2ABZ | 160 | 1871 | train | 2031 | 7,9 % |
| 2AJF | 242 | 2288 | train | 2530 | 9,6 % |
| 2AYO | 73 | 2457 | train | 2530 | 2,9 % |
| 2B42 | 180 | 2350 | train | 2530 | 7,1 % |
| 2BTF | 232 | 2091 | train | 2323 | 10,0 % |
| 2C0L | 177 | 1320 | eval | 1497 | 11,8 % |
| 2CFH | 206 | 2324 | train | 2530 | 8,1 % |
| 2FJU | 309 | 2220 | train | 2529 | 12,2 % |
| 2G77 | 112 | 2418 | eval | 2530 | 4,4 % |
| 2GAF | 152 | 2370 | train | 2522 | 6,0 % |
| 2GTP | 195 | 905 | train | 1100 | 17,7 % |
| 2H7V | 10 | 2231 | train | 2241 | 0,4 % |
| 2HLE | 117 | 2413 | train | 2530 | 4,6 % |
| 2HQS | 238 | 2292 | train | 2530 | 9,4 % |
| 2HRK | 10 | 2392 | train | 2402 | 0,4 % |
| 2I25 | 284 | 2246 | train | 2530 | 11,2 % |
| 2I9B | 89 | 2441 | train | 2530 | 3,5 % |
| 2IDO | 73 | 2457 | train | 2530 | 2,9 % |
| 2J0T | 289 | 1730 | train | 2019 | 14,3 % |
| 2J7P | 22 | 2508 | eval | 2530 | 0,9 % |
| 2NZ8 | 44 | 1146 | train | 1190 | 3,7 % |
| 2O3B | 39 | 2491 | train | 2530 | 1,5 % |
| 2O8V | 226 | 2304 | train | 2530 | 8,9 % |
| 2OOB | 319 | 2211 | train | 2530 | 12,6 % |
| 2OT3 | 17 | 2513 | train | 2530 | 0,7 % |
| 2OUL | 298 | 1349 | eval | 1647 | 18,1 % |
| 2PCC | 131 | 2399 | train | 2530 | 5,2 % |
| 2SIC | 278 | 2252 | train | 2530 | 11,0 % |
| 2UUY | 300 | 2230 | train | 2530 | 11,9 % |
| 2VDB | 264 | 1382 | train | 1646 | 16,0 % |
| 2X9A | 94 | 2436 | train | 2530 | 3,7 % |
| 3A4S | 118 | 2390 | eval | 2508 | 4,7 % |
| 3AAD | 12 | 2518 | train | 2530 | 0,5 % |
| 3BIW | 35 | 2348 | train | 2383 | 1,5 % |
| 3BX7 | 19 | 1972 | train | 1991 | 1,0 % |
| 3CPH | 59 | 1984 | train | 2043 | 2,9 % |
| 3D5S | 258 | 2272 | train | 2530 | 10,2 % |
| 3DAW | 135 | 2055 | train | 2190 | 6,2 % |
| 3F1P | 104 | 2261 | eval | 2365 | 4,4 % |
| 3FN1 | 24 | 2506 | train | 2530 | 0,9 % |
| 3H2V | 159 | 2371 | eval | 2530 | 6,3 % |
| 3PC8 | 321 | 2209 | train | 2530 | 12,7 % |
| 3S9D | 49 | 2481 | train | 2530 | 1,9 % |
| 3SGQ | 92 | 2438 | train | 2530 | 3,6 % |
| 3VLB | 268 | 2262 | eval | 2530 | 10,6 % |
| 4CPA | 295 | 2235 | train | 2530 | 11,7 % |
| 4FZA | 38 | 2492 | train | 2530 | 1,5 % |
| 4H03 | 275 | 2255 | train | 2530 | 10,9 % |
| 4IZ7 | 18 | 1068 | eval | 1086 | 1,7 % |
| 4M76 | 346 | 2184 | train | 2530 | 13,7 % |
| 7CEI | 210 | 2320 | train | 2530 | 8,3 % |
| BAAD | 198 | 2332 | train | 2530 | 7,8 % |
| BOYV | 257 | 2273 | train | 2530 | 10,2 % |

Table S9 Training and evaluation set composition – fold8

| fold8 | fnat ≥ 0.3 | fnat < 0.3 | set | total | % good models |
| --- | --- | --- | --- | --- | --- |
| 1ACB | 128 | 1908 | train | 2036 | 6,3 % |
| 1ATN | 82 | 2448 | train | 2530 | 3,2 % |
| 1AVX | 153 | 1526 | train | 1679 | 9,1 % |
| 1AY7 | 276 | 2254 | train | 2530 | 10,9 % |
| 1B6C | 10 | 2316 | train | 2326 | 0,4 % |
| 1BKD | 13 | 2517 | train | 2530 | 0,5 % |
| 1BUH | 213 | 135 | eval | 348 | 61,2 % |
| 1CLV | 78 | 2452 | train | 2530 | 3,1 % |
| 1D6R | 117 | 2413 | train | 2530 | 4,6 % |
| 1DFJ | 241 | 684 | train | 925 | 26,1 % |
| 1E6E | 312 | 1901 | train | 2213 | 14,1 % |
| 1E96 | 289 | 1190 | train | 1479 | 19,5 % |
| 1EAW | 110 | 472 | train | 582 | 18,9 % |
| 1EFN | 112 | 2418 | train | 2530 | 4,4 % |
| 1EWY | 201 | 2329 | train | 2530 | 7,9 % |
| 1F34 | 176 | 591 | train | 767 | 22,9 % |
| 1FC2 | 183 | 1010 | eval | 1193 | 15,3 % |
| 1FFW | 268 | 2262 | train | 2530 | 10,6 % |
| 1FLE | 140 | 2390 | train | 2530 | 5,5 % |
| 1FQ1 | 135 | 2395 | train | 2530 | 5,3 % |
| 1FQJ | 267 | 2263 | eval | 2530 | 10,6 % |
| 1GCQ | 265 | 2265 | eval | 2530 | 10,5 % |
| 1GHQ | 145 | 2385 | train | 2530 | 5,7 % |
| 1GL1 | 116 | 2414 | train | 2530 | 4,6 % |
| 1GLA | 126 | 2404 | train | 2530 | 5,0 % |
| 1GPW | 139 | 2391 | train | 2530 | 5,5 % |
| 1GRN | 162 | 1579 | train | 1741 | 9,3 % |
| 1GXD | 249 | 2281 | train | 2530 | 9,8 % |
| 1H9D | 127 | 1806 | eval | 1933 | 6,6 % |
| 1HE1 | 209 | 1929 | eval | 2138 | 9,8 % |
| 1HE8 | 145 | 2385 | train | 2530 | 5,7 % |
| 1I2M | 12 | 2518 | train | 2530 | 0,5 % |
| 1J2J | 278 | 2252 | train | 2530 | 11,0 % |
| 1JIW | 24 | 2240 | train | 2264 | 1,1 % |
| 1JK9 | 273 | 2257 | train | 2530 | 10,8 % |
| 1JTD | 164 | 2366 | train | 2530 | 6,5 % |
| 1JTG | 125 | 2405 | eval | 2530 | 4,9 % |
| 1KAC | 253 | 2277 | train | 2530 | 10,0 % |
| 1KTZ | 129 | 2401 | eval | 2530 | 5,1 % |
| 1KXP | 10 | 2216 | train | 2226 | 0,4 % |
| 1KXQ | 256 | 2274 | train | 2530 | 10,1 % |
| 1LFD | 116 | 2414 | train | 2530 | 4,6 % |
| 1M10 | 173 | 2357 | train | 2530 | 6,8 % |
| 1MAH | 317 | 2213 | eval | 2530 | 12,5 % |
| 1MQ8 | 285 | 2245 | train | 2530 | 11,3 % |
| 1NW9 | 52 | 2478 | eval | 2530 | 2,1 % |
| 1OC0 | 124 | 1610 | eval | 1734 | 7,2 % |
| 1PVH | 9 | 2271 | train | 2280 | 0,4 % |
| 1PXV | 205 | 2140 | train | 2345 | 8,7 % |
| 1QA9 | 292 | 1277 | train | 1569 | 18,6 % |
| 1R0R | 278 | 2252 | train | 2530 | 11,0 % |
| 1R6Q | 233 | 2297 | train | 2530 | 9,2 % |
| 1R8S | 10 | 1576 | train | 1586 | 0,6 % |
| 1RKE | 16 | 1321 | eval | 1337 | 1,2 % |
| 1S1Q | 235 | 2295 | train | 2530 | 9,3 % |
| 1SBB | 160 | 1070 | train | 1230 | 13,0 % |
| 1SYX | 278 | 2252 | train | 2530 | 11,0 % |
| 1T6B | 28 | 1299 | train | 1327 | 2,1 % |
| 1TMQ | 298 | 2232 | train | 2530 | 11,8 % |
| 1UDI | 114 | 244 | eval | 358 | 31,8 % |
| 1US7 | 87 | 950 | train | 1037 | 8,4 % |
| 1WQ1 | 33 | 2497 | train | 2530 | 1,3 % |
| 1XD3 | 11 | 100 | train | 111 | 9,9 % |
| 1Y64 | 10 | 1416 | train | 1426 | 0,7 % |
| 1YVB | 171 | 2359 | eval | 2530 | 6,8 % |
| 1Z0K | 230 | 2300 | eval | 2530 | 9,1 % |
| 1Z5Y | 168 | 2362 | train | 2530 | 6,6 % |
| 1ZHH | 291 | 2239 | train | 2530 | 11,5 % |
| 1ZHI | 168 | 2362 | train | 2530 | 6,6 % |
| 1ZLI | 49 | 1354 | train | 1403 | 3,5 % |
| 1ZM4 | 252 | 1118 | eval | 1370 | 18,4 % |
| 2A1A | 151 | 2379 | train | 2530 | 6,0 % |
| 2A5T | 40 | 2490 | train | 2530 | 1,6 % |
| 2A9K | 56 | 2474 | train | 2530 | 2,2 % |
| 2ABZ | 156 | 1857 | eval | 2013 | 7,7 % |
| 2AJF | 245 | 2285 | train | 2530 | 9,7 % |
| 2AYO | 198 | 2332 | eval | 2530 | 7,8 % |
| 2B42 | 226 | 2304 | train | 2530 | 8,9 % |
| 2BTF | 233 | 2109 | train | 2342 | 9,9 % |
| 2C0L | 185 | 1316 | train | 1501 | 12,3 % |
| 2CFH | 208 | 2322 | train | 2530 | 8,2 % |
| 2FJU | 310 | 2220 | train | 2530 | 12,3 % |
| 2G77 | 130 | 2400 | train | 2530 | 5,1 % |
| 2GAF | 140 | 2390 | train | 2530 | 5,5 % |
| 2GTP | 194 | 940 | train | 1134 | 17,1 % |
| 2H7V | 10 | 2223 | train | 2233 | 0,4 % |
| 2HLE | 108 | 2422 | train | 2530 | 4,3 % |
| 2HQS | 250 | 2280 | eval | 2530 | 9,9 % |
| 2HRK | 11 | 2389 | eval | 2400 | 0,5 % |
| 2I25 | 285 | 2245 | train | 2530 | 11,3 % |
| 2I9B | 98 | 2432 | train | 2530 | 3,9 % |
| 2IDO | 131 | 2399 | eval | 2530 | 5,2 % |
| 2J0T | 295 | 1732 | train | 2027 | 14,6 % |
| 2J7P | 89 | 2441 | train | 2530 | 3,5 % |
| 2NZ8 | 29 | 1183 | train | 1212 | 2,4 % |
| 2O3B | 24 | 2506 | train | 2530 | 0,9 % |
| 2O8V | 245 | 2285 | train | 2530 | 9,7 % |
| 2OOB | 320 | 2210 | train | 2530 | 12,6 % |
| 2OT3 | 16 | 2514 | train | 2530 | 0,6 % |
| 2OUL | 298 | 1381 | train | 1679 | 17,7 % |
| 2PCC | 134 | 2396 | train | 2530 | 5,3 % |
| 2SIC | 279 | 2251 | train | 2530 | 11,0 % |
| 2UUY | 328 | 2202 | train | 2530 | 13,0 % |
| 2VDB | 276 | 1358 | train | 1634 | 16,9 % |
| 2X9A | 254 | 2276 | train | 2530 | 10,0 % |
| 3A4S | 113 | 2417 | train | 2530 | 4,5 % |
| 3AAD | 11 | 2519 | train | 2530 | 0,4 % |
| 3BIW | 33 | 2343 | train | 2376 | 1,4 % |
| 3BX7 | 74 | 1939 | train | 2013 | 3,7 % |
| 3CPH | 85 | 1992 | train | 2077 | 4,1 % |
| 3D5S | 259 | 2271 | train | 2530 | 10,2 % |
| 3DAW | 159 | 1973 | train | 2132 | 7,5 % |
| 3F1P | 103 | 2274 | train | 2377 | 4,3 % |
| 3FN1 | 16 | 2514 | train | 2530 | 0,6 % |
| 3H2V | 166 | 2364 | train | 2530 | 6,6 % |
| 3PC8 | 323 | 2207 | eval | 2530 | 12,8 % |
| 3S9D | 43 | 2487 | train | 2530 | 1,7 % |
| 3SGQ | 204 | 2326 | eval | 2530 | 8,1 % |
| 3VLB | 264 | 2266 | train | 2530 | 10,4 % |
| 4CPA | 291 | 2239 | train | 2530 | 11,5 % |
| 4FZA | 140 | 2390 | eval | 2530 | 5,5 % |
| 4H03 | 274 | 2256 | eval | 2530 | 10,8 % |
| 4IZ7 | 12 | 1076 | train | 1088 | 1,1 % |
| 4M76 | 344 | 2186 | train | 2530 | 13,6 % |
| 7CEI | 218 | 2312 | train | 2530 | 8,6 % |
| BAAD | 242 | 2288 | train | 2530 | 9,6 % |
| BOYV | 251 | 2279 | train | 2530 | 9,9 % |

Table S10 Training and evaluation set composition – fold9

| fold9 | fnat ≥ 0.3 | fnat < 0.3 | set | total | % good models |
| --- | --- | --- | --- | --- | --- |
| 1ACB | 235 | 1912 | train | 2147 | 10,9 % |
| 1ATN | 89 | 2441 | eval | 2530 | 3,5 % |
| 1AVX | 161 | 1523 | train | 1684 | 9,6 % |
| 1AY7 | 277 | 2253 | train | 2530 | 10,9 % |
| 1B6C | 10 | 2316 | train | 2326 | 0,4 % |
| 1BKD | 26 | 2504 | train | 2530 | 1,0 % |
| 1BUH | 210 | 141 | train | 351 | 59,8 % |
| 1CLV | 119 | 2411 | train | 2530 | 4,7 % |
| 1D6R | 115 | 2415 | train | 2530 | 4,5 % |
| 1DFJ | 254 | 682 | train | 936 | 27,1 % |
| 1E6E | 314 | 1901 | train | 2215 | 14,2 % |
| 1E96 | 294 | 1206 | eval | 1500 | 19,6 % |
| 1EAW | 116 | 469 | train | 585 | 19,8 % |
| 1EFN | 124 | 2406 | eval | 2530 | 4,9 % |
| 1EWY | 201 | 2329 | train | 2530 | 7,9 % |
| 1F34 | 312 | 484 | eval | 796 | 39,2 % |
| 1FC2 | 196 | 989 | train | 1185 | 16,5 % |
| 1FFW | 275 | 2255 | eval | 2530 | 10,9 % |
| 1FLE | 196 | 2334 | eval | 2530 | 7,7 % |
| 1FQ1 | 139 | 2391 | train | 2530 | 5,5 % |
| 1FQJ | 273 | 2257 | train | 2530 | 10,8 % |
| 1GCQ | 270 | 2260 | train | 2530 | 10,7 % |
| 1GHQ | 138 | 2392 | train | 2530 | 5,5 % |
| 1GL1 | 108 | 2422 | train | 2530 | 4,3 % |
| 1GLA | 244 | 2286 | eval | 2530 | 9,6 % |
| 1GPW | 136 | 2394 | eval | 2530 | 5,4 % |
| 1GRN | 247 | 1452 | train | 1699 | 14,5 % |
| 1GXD | 237 | 2293 | train | 2530 | 9,4 % |
| 1H9D | 146 | 1811 | train | 1957 | 7,5 % |
| 1HE1 | 181 | 1971 | train | 2152 | 8,4 % |
| 1HE8 | 142 | 2388 | train | 2530 | 5,6 % |
| 1I2M | 68 | 2462 | train | 2530 | 2,7 % |
| 1J2J | 300 | 2230 | train | 2530 | 11,9 % |
| 1JIW | 28 | 2236 | train | 2264 | 1,2 % |
| 1JK9 | 284 | 2246 | eval | 2530 | 11,2 % |
| 1JTD | 167 | 2236 | train | 2403 | 6,9 % |
| 1JTG | 128 | 2312 | train | 2440 | 5,2 % |
| 1KAC | 257 | 2273 | train | 2530 | 10,2 % |
| 1KTZ | 135 | 2395 | train | 2530 | 5,3 % |
| 1KXP | 10 | 2215 | train | 2225 | 0,4 % |
| 1KXQ | 247 | 2283 | train | 2530 | 9,8 % |
| 1LFD | 72 | 2458 | train | 2530 | 2,8 % |
| 1M10 | 166 | 2364 | train | 2530 | 6,6 % |
| 1MAH | 316 | 2214 | train | 2530 | 12,5 % |
| 1MQ8 | 284 | 2246 | train | 2530 | 11,2 % |
| 1NW9 | 94 | 2436 | train | 2530 | 3,7 % |
| 1OC0 | 143 | 1585 | train | 1728 | 8,3 % |
| 1PVH | 9 | 2271 | train | 2280 | 0,4 % |
| 1PXV | 195 | 2150 | train | 2345 | 8,3 % |
| 1QA9 | 291 | 1244 | train | 1535 | 19,0 % |
| 1R0R | 276 | 2254 | train | 2530 | 10,9 % |
| 1R6Q | 236 | 2294 | train | 2530 | 9,3 % |
| 1R8S | 10 | 1809 | train | 1819 | 0,5 % |
| 1RKE | 22 | 1596 | train | 1618 | 1,4 % |
| 1S1Q | 234 | 2296 | train | 2530 | 9,2 % |
| 1SBB | 132 | 1008 | train | 1140 | 11,6 % |
| 1SYX | 301 | 2229 | train | 2530 | 11,9 % |
| 1T6B | 0 | 1273 | eval | 1273 | 0,0 % |
| 1TMQ | 297 | 2233 | train | 2530 | 11,7 % |
| 1UDI | 166 | 203 | train | 369 | 45,0 % |
| 1US7 | 92 | 883 | train | 975 | 9,4 % |
| 1WQ1 | 30 | 2500 | train | 2530 | 1,2 % |
| 1XD3 | 13 | 84 | train | 97 | 13,4 % |
| 1Y64 | 10 | 1895 | train | 1905 | 0,5 % |
| 1YVB | 171 | 2359 | train | 2530 | 6,8 % |
| 1Z0K | 230 | 2300 | train | 2530 | 9,1 % |
| 1Z5Y | 198 | 2332 | train | 2530 | 7,8 % |
| 1ZHH | 281 | 2249 | train | 2530 | 11,1 % |
| 1ZHI | 158 | 2372 | eval | 2530 | 6,2 % |
| 1ZLI | 79 | 1434 | train | 1513 | 5,2 % |
| 1ZM4 | 242 | 1134 | train | 1376 | 17,6 % |
| 2A1A | 158 | 2372 | train | 2530 | 6,2 % |
| 2A5T | 131 | 2399 | train | 2530 | 5,2 % |
| 2A9K | 54 | 2476 | train | 2530 | 2,1 % |
| 2ABZ | 164 | 1831 | train | 1995 | 8,2 % |
| 2AJF | 253 | 2277 | eval | 2530 | 10,0 % |
| 2AYO | 200 | 2330 | train | 2530 | 7,9 % |
| 2B42 | 224 | 2306 | train | 2530 | 8,9 % |
| 2BTF | 293 | 2062 | eval | 2355 | 12,4 % |
| 2C0L | 188 | 1327 | train | 1515 | 12,4 % |
| 2CFH | 176 | 2354 | train | 2530 | 7,0 % |
| 2FJU | 246 | 1688 | train | 1934 | 12,7 % |
| 2G77 | 145 | 2385 | train | 2530 | 5,7 % |
| 2GAF | 195 | 2335 | eval | 2530 | 7,7 % |
| 2GTP | 203 | 905 | train | 1108 | 18,3 % |
| 2H7V | 10 | 2195 | eval | 2205 | 0,5 % |
| 2HLE | 176 | 2354 | eval | 2530 | 7,0 % |
| 2HQS | 271 | 2259 | train | 2530 | 10,7 % |
| 2HRK | 10 | 2390 | train | 2400 | 0,4 % |
| 2I25 | 305 | 2225 | eval | 2530 | 12,1 % |
| 2I9B | 86 | 2444 | eval | 2530 | 3,4 % |
| 2IDO | 162 | 2368 | train | 2530 | 6,4 % |
| 2J0T | 303 | 1667 | train | 1970 | 15,4 % |
| 2J7P | 89 | 2441 | train | 2530 | 3,5 % |
| 2NZ8 | 56 | 1185 | train | 1241 | 4,5 % |
| 2O3B | 24 | 2506 | train | 2530 | 0,9 % |
| 2O8V | 246 | 2284 | eval | 2530 | 9,7 % |
| 2OOB | 323 | 2207 | train | 2530 | 12,8 % |
| 2OT3 | 36 | 2494 | train | 2530 | 1,4 % |
| 2OUL | 297 | 1373 | train | 1670 | 17,8 % |
| 2PCC | 132 | 2398 | train | 2530 | 5,2 % |
| 2SIC | 294 | 2236 | train | 2530 | 11,6 % |
| 2UUY | 295 | 2235 | train | 2530 | 11,7 % |
| 2VDB | 281 | 1390 | train | 1671 | 16,8 % |
| 2X9A | 260 | 2270 | eval | 2530 | 10,3 % |
| 3A4S | 119 | 2410 | train | 2529 | 4,7 % |
| 3AAD | 23 | 2507 | train | 2530 | 0,9 % |
| 3BIW | 31 | 2243 | train | 2274 | 1,4 % |
| 3BX7 | 187 | 1758 | train | 1945 | 9,6 % |
| 3CPH | 90 | 1979 | train | 2069 | 4,3 % |
| 3D5S | 272 | 2258 | train | 2530 | 10,8 % |
| 3DAW | 238 | 1940 | eval | 2178 | 10,9 % |
| 3F1P | 127 | 2260 | train | 2387 | 5,3 % |
| 3FN1 | 13 | 2517 | eval | 2530 | 0,5 % |
| 3H2V | 152 | 2378 | train | 2530 | 6,0 % |
| 3PC8 | 325 | 2205 | train | 2530 | 12,8 % |
| 3S9D | 43 | 2487 | train | 2530 | 1,7 % |
| 3SGQ | 275 | 2255 | train | 2530 | 10,9 % |
| 3VLB | 264 | 2266 | train | 2530 | 10,4 % |
| 4CPA | 295 | 2235 | eval | 2530 | 11,7 % |
| 4FZA | 191 | 2339 | train | 2530 | 7,5 % |
| 4H03 | 295 | 2235 | train | 2530 | 11,7 % |
| 4IZ7 | 12 | 1145 | train | 1157 | 1,0 % |
| 4M76 | 345 | 2185 | train | 2530 | 13,6 % |
| 7CEI | 228 | 2302 | eval | 2530 | 9,0 % |
| BAAD | 236 | 2294 | train | 2530 | 9,3 % |
| BOYV | 250 | 2280 | eval | 2530 | 9,9 % |

Table S11 Training and evaluation set composition – fold10

| fold10 | fnat ≥ 0.3 | fnat < 0.3 | set | total | % good models |
| --- | --- | --- | --- | --- | --- |
| 1ACB | 765 | 1764 | train | 2529 | 30,2 % |
| 1ATN | 258 | 2272 | train | 2530 | 10,2 % |
| 1AVX | 176 | 1907 | eval | 2083 | 8,4 % |
| 1AY7 | 329 | 2201 | train | 2530 | 13,0 % |
| 1B6C | 12 | 1838 | train | 1850 | 0,6 % |
| 1BKD | 25 | 2505 | eval | 2530 | 1,0 % |
| 1BUH | 220 | 1296 | train | 1516 | 14,5 % |
| 1CLV | 104 | 2426 | train | 2530 | 4,1 % |
| 1D6R | 116 | 2414 | eval | 2530 | 4,6 % |
| 1DFJ | 254 | 1356 | train | 1610 | 15,8 % |
| 1E6E | 313 | 1970 | train | 2283 | 13,7 % |
| 1E96 | 307 | 1456 | train | 1763 | 17,4 % |
| 1EAW | 152 | 2010 | eval | 2162 | 7,0 % |
| 1EFN | 164 | 2366 | train | 2530 | 6,5 % |
| 1EWY | 201 | 2328 | train | 2529 | 7,9 % |
| 1F34 | 312 | 732 | train | 1044 | 29,9 % |
| 1FC2 | 226 | 1746 | train | 1972 | 11,5 % |
| 1FFW | 284 | 2246 | train | 2530 | 11,2 % |
| 1FLE | 217 | 2313 | train | 2530 | 8,6 % |
| 1FQ1 | 292 | 2238 | eval | 2530 | 11,5 % |
| 1FQJ | 304 | 2226 | train | 2530 | 12,0 % |
| 1GCQ | 269 | 2261 | train | 2530 | 10,6 % |
| 1GHQ | 597 | 1933 | train | 2530 | 23,6 % |
| 1GL1 | 97 | 2433 | eval | 2530 | 3,8 % |
| 1GLA | 325 | 2205 | train | 2530 | 12,8 % |
| 1GPW | 161 | 2369 | train | 2530 | 6,4 % |
| 1GRN | 301 | 1718 | train | 2019 | 14,9 % |
| 1GXD | 242 | 2288 | train | 2530 | 9,6 % |
| 1H9D | 164 | 2210 | train | 2374 | 6,9 % |
| 1HE1 | 183 | 2221 | train | 2404 | 7,6 % |
| 1HE8 | 748 | 1782 | train | 2530 | 29,6 % |
| 1I2M | 81 | 2449 | train | 2530 | 3,2 % |
| 1J2J | 339 | 2191 | eval | 2530 | 13,4 % |
| 1JIW | 18 | 2141 | train | 2159 | 0,8 % |
| 1JK9 | 302 | 2228 | train | 2530 | 11,9 % |
| 1JTD | 177 | 2060 | train | 2237 | 7,9 % |
| 1JTG | 182 | 2303 | train | 2485 | 7,3 % |
| 1KAC | 250 | 2280 | train | 2530 | 9,9 % |
| 1KTZ | 134 | 2396 | train | 2530 | 5,3 % |
| 1KXP | 10 | 2003 | eval | 2013 | 0,5 % |
| 1KXQ | 247 | 2283 | train | 2530 | 9,8 % |
| 1LFD | 393 | 2137 | train | 2530 | 15,5 % |
| 1M10 | 166 | 2364 | train | 2530 | 6,6 % |
| 1MAH | 314 | 2216 | train | 2530 | 12,4 % |
| 1MQ8 | 282 | 2248 | train | 2530 | 11,1 % |
| 1NW9 | 117 | 2413 | train | 2530 | 4,6 % |
| 1OC0 | 147 | 2257 | train | 2404 | 6,1 % |
| 1PVH | 199 | 2035 | eval | 2234 | 8,9 % |
| 1PXV | 211 | 2244 | train | 2455 | 8,6 % |
| 1QA9 | 293 | 1442 | train | 1735 | 16,9 % |
| 1R0R | 276 | 2254 | train | 2530 | 10,9 % |
| 1R6Q | 250 | 2280 | eval | 2530 | 9,9 % |
| 1R8S | 10 | 2520 | train | 2530 | 0,4 % |
| 1RKE | 14 | 2516 | train | 2530 | 0,6 % |
| 1S1Q | 239 | 2289 | eval | 2528 | 9,5 % |
| 1SBB | 209 | 2126 | train | 2335 | 9,0 % |
| 1SYX | 331 | 2199 | train | 2530 | 13,1 % |
| 1T6B | 2 | 1506 | train | 1508 | 0,1 % |
| 1TMQ | 297 | 2233 | eval | 2530 | 11,7 % |
| 1US7 | 179 | 803 | train | 982 | 18,2 % |
| 1WQ1 | 61 | 2469 | eval | 2530 | 2,4 % |
| 1XD3 | 11 | 246 | train | 257 | 4,3 % |
| 1Y64 | 15 | 2515 | train | 2530 | 0,6 % |
| 1YVB | 166 | 2364 | train | 2530 | 6,6 % |
| 1Z0K | 245 | 2285 | train | 2530 | 9,7 % |
| 1Z5Y | 255 | 2275 | train | 2530 | 10,1 % |
| 1ZHH | 283 | 2247 | train | 2530 | 11,2 % |
| 1ZHI | 184 | 2346 | train | 2530 | 7,3 % |
| 1ZLI | 49 | 2273 | eval | 2322 | 2,1 % |
| 1ZM4 | 244 | 1555 | train | 1799 | 13,6 % |
| 2A1A | 199 | 2329 | train | 2528 | 7,9 % |
| 2A5T | 423 | 2107 | train | 2530 | 16,7 % |
| 2A9K | 55 | 2475 | train | 2530 | 2,2 % |
| 2ABZ | 266 | 1917 | train | 2183 | 12,2 % |
| 2AJF | 254 | 2276 | train | 2530 | 10,0 % |
| 2AYO | 200 | 2330 | train | 2530 | 7,9 % |
| 2B42 | 224 | 2304 | train | 2528 | 8,9 % |
| 2BTF | 300 | 2103 | train | 2403 | 12,5 % |
| 2C0L | 212 | 1979 | train | 2191 | 9,7 % |
| 2CFH | 213 | 2317 | eval | 2530 | 8,4 % |
| 2FJU | 345 | 1604 | train | 1949 | 17,7 % |
| 2G77 | 174 | 2356 | train | 2530 | 6,9 % |
| 2GAF | 231 | 1614 | train | 1845 | 12,5 % |
| 2GTP | 202 | 1894 | eval | 2096 | 9,6 % |
| 2H7V | 15 | 1337 | train | 1352 | 1,1 % |
| 2HLE | 204 | 2326 | train | 2530 | 8,1 % |
| 2HQS | 274 | 2256 | train | 2530 | 10,8 % |
| 2HRK | 10 | 1644 | train | 1654 | 0,6 % |
| 2I25 | 305 | 2225 | train | 2530 | 12,1 % |
| 2I9B | 106 | 2424 | train | 2530 | 4,2 % |
| 2IDO | 171 | 2359 | train | 2530 | 6,8 % |
| 2J0T | 303 | 1849 | eval | 2152 | 14,1 % |
| 2J7P | 92 | 2438 | train | 2530 | 3,6 % |
| 2NZ8 | 47 | 2158 | train | 2205 | 2,1 % |
| 2O3B | 37 | 2493 | eval | 2530 | 1,5 % |
| 2O8V | 842 | 1688 | train | 2530 | 33,3 % |
| 2OOB | 387 | 2140 | eval | 2527 | 15,3 % |
| 2OT3 | 72 | 2458 | eval | 2530 | 2,8 % |
| 2OUL | 297 | 1577 | train | 1874 | 15,8 % |
| 2PCC | 134 | 2396 | train | 2530 | 5,3 % |
| 2SIC | 298 | 2232 | eval | 2530 | 11,8 % |
| 2UUY | 322 | 2208 | train | 2530 | 12,7 % |
| 2VDB | 281 | 1653 | eval | 1934 | 14,5 % |
| 2X9A | 259 | 2271 | train | 2530 | 10,2 % |
| 3A4S | 566 | 1720 | train | 2286 | 24,8 % |
| 3AAD | 99 | 2431 | eval | 2530 | 3,9 % |
| 3BIW | 10 | 897 | train | 907 | 1,1 % |
| 3BX7 | 209 | 2114 | train | 2323 | 9,0 % |
| 3CPH | 139 | 2259 | eval | 2398 | 5,8 % |
| 3D5S | 272 | 2258 | eval | 2530 | 10,8 % |
| 3DAW | 240 | 2093 | train | 2333 | 10,3 % |
| 3F1P | 145 | 2357 | train | 2502 | 5,8 % |
| 3FN1 | 16 | 2514 | train | 2530 | 0,6 % |
| 3H2V | 153 | 2377 | train | 2530 | 6,0 % |
| 3PC8 | 338 | 2192 | train | 2530 | 13,4 % |
| 3S9D | 61 | 2469 | train | 2530 | 2,4 % |
| 3SGQ | 284 | 2246 | train | 2530 | 11,2 % |
| 3VLB | 264 | 2266 | train | 2530 | 10,4 % |
| 4CPA | 312 | 2218 | train | 2530 | 12,3 % |
| 4FZA | 196 | 2334 | train | 2530 | 7,7 % |
| 4H03 | 322 | 2208 | train | 2530 | 12,7 % |
| 4IZ7 | 41 | 2489 | train | 2530 | 1,6 % |
| 4M76 | 343 | 2187 | train | 2530 | 13,6 % |
| 7CEI | 275 | 2255 | train | 2530 | 10,9 % |
| BAAD | 255 | 2275 | train | 2530 | 10,1 % |
| BOYV | 269 | 2261 | train | 2530 | 10,6 % |

Table S12: CAPRI quality criteria automatically computed by DeepRank-GNN upon provision of a reference structure

| CAPRI quality criteria | Definition |
| --- | --- |
| interface RMSD (iRMSD) | RMSD between superimposed interface residues |
| ligand-RMSD (lRMSD) | RMSD between chains B after superimposition of chains A |
| fraction of native contacts (*f_nat_*) | the fraction of reference interface contacts preserved in the interface of the docking model; the interface is defined as any pair of heavy atoms from two chains within 5Å of each other. |
| dockQ | (Basu and Wallner, 2016) |
| binary class | $0:irmsd\geq4 Å, 1:irmsd<4Å$ |
| capri irmsd classes | $1:irmsd<1Å, 2:irmsd<2Å,3:irmsd<4Å, 4:irmsd<6Å, 0:irmsd\geq6 Å$ |

**Figure S1** (A) Correlation between iRMSD and fnat values, (B) superimposition of a docking model (green and cyan) associated to a fnat > 0.6 and an iRMSD > 6A on the reference structure (1JTG - grey).

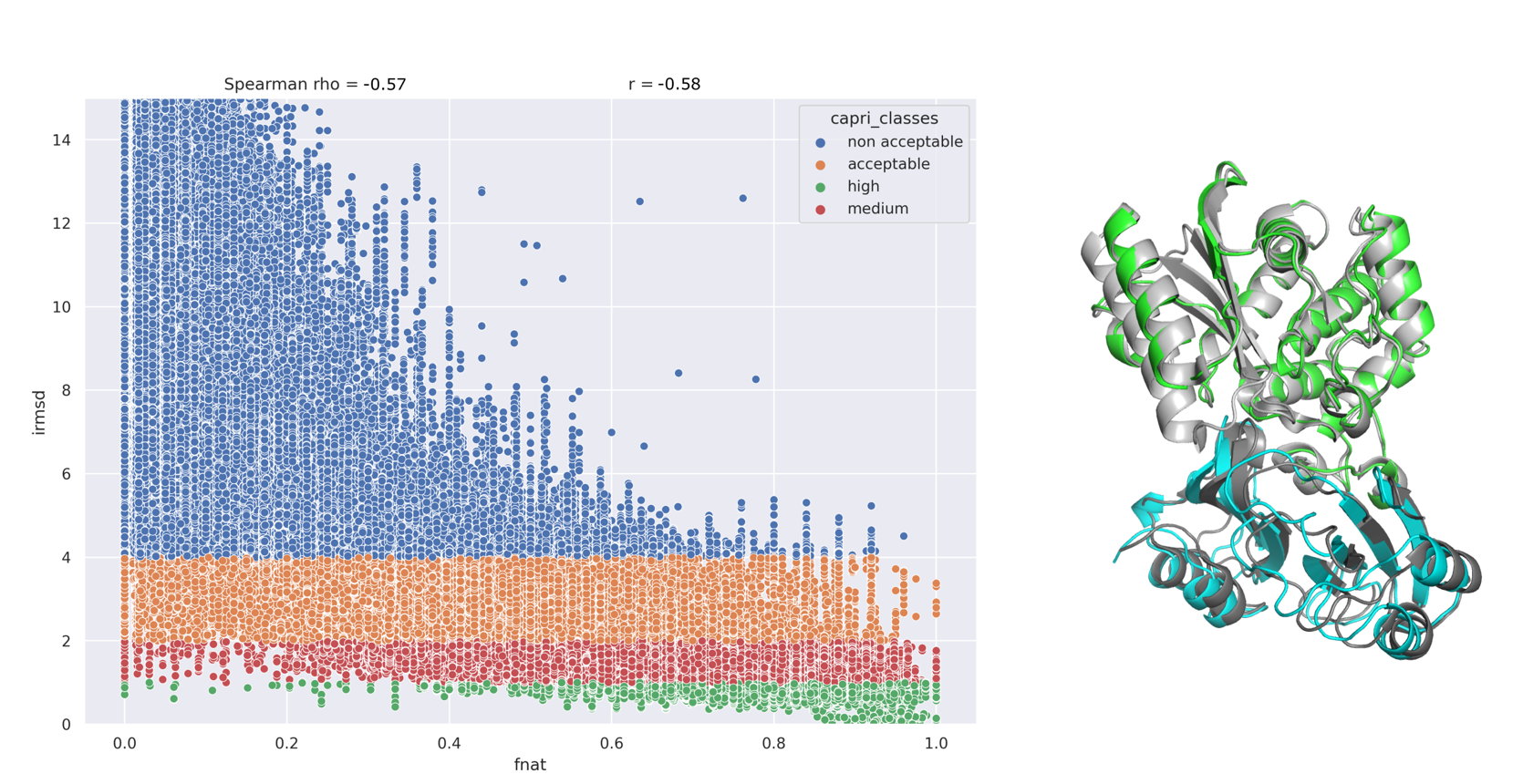

**Figure S2** DeepRank-GNN diagram. The graphs generation module of DeepRank-GNN requires PDB files, features and target specifications as input data. The target values can be either provided by the user or automatically computed in a docking benchmark scenario upon provision of reference structures. Generated graphs are stored in HDF5 format for memory and I/O optimization. Graphs are fed to the training module that additionally requires a defined GNN architecture and hyperparameters. The user can save all generated models or save only the last one, intermediate models, or the best one based on the loss value on the evaluation set. The trained model(s) can be further applied to test sets and unlabelled graphs for independent validation and prediction.

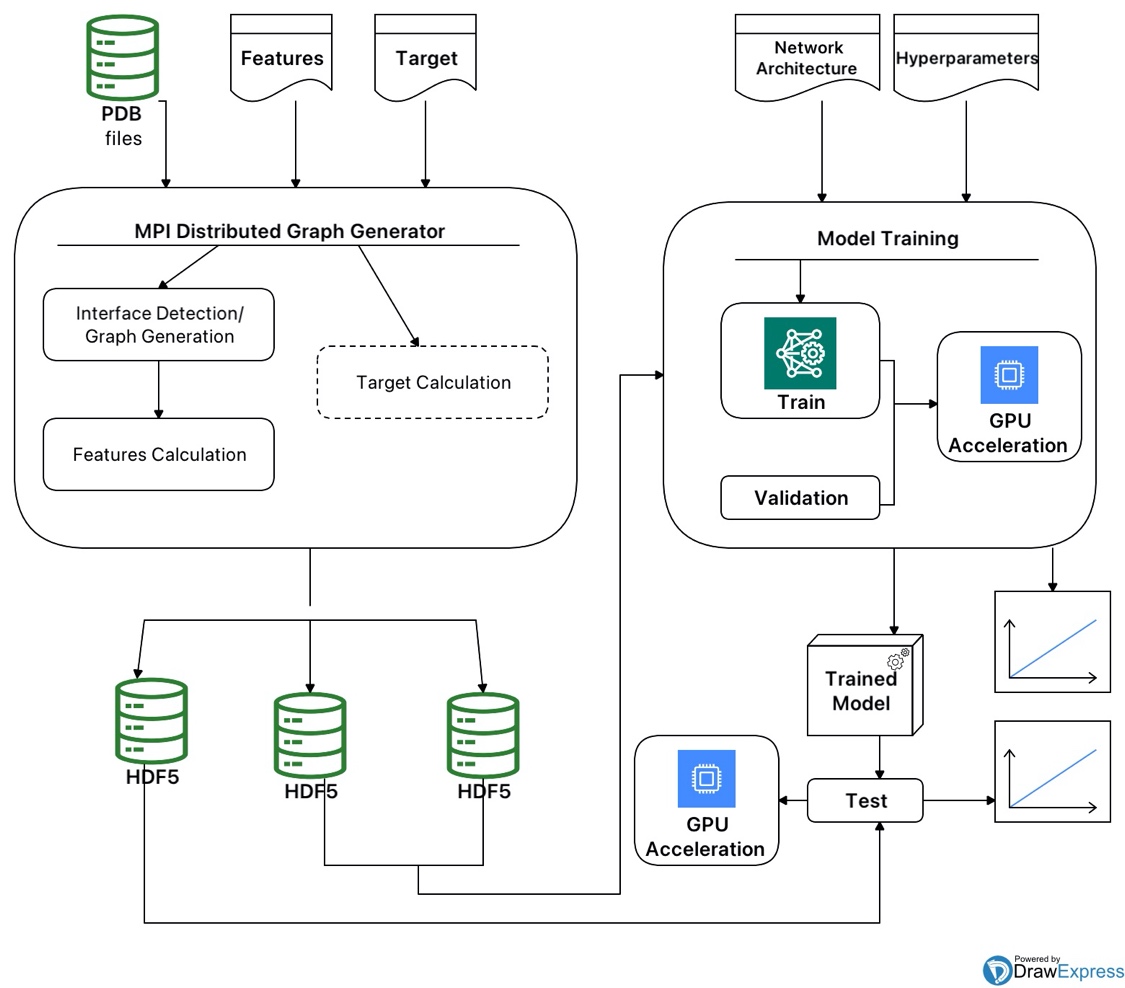

**Figure S3** Evolution of the loss value obtained on the training and the evaluation sets over 20 epochs. Results are shown for the 10 -folds of the cross validation considering PSSM information or not.

**Figure S4**: Average Receiver operating characteristic curves (ROC) obtained with the models retained for each DeepRank-GNN fold and HADDOCK score. A true positive case corresponds to a complex with fnat ≥ 0.3 correctly predicted. The number of True Positive Rate value is averaged over the number of complexes in the test dataset. The dashed line represents a random classifier

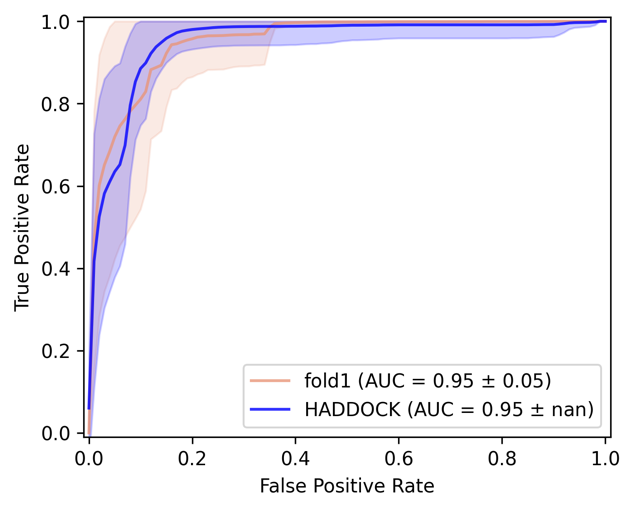

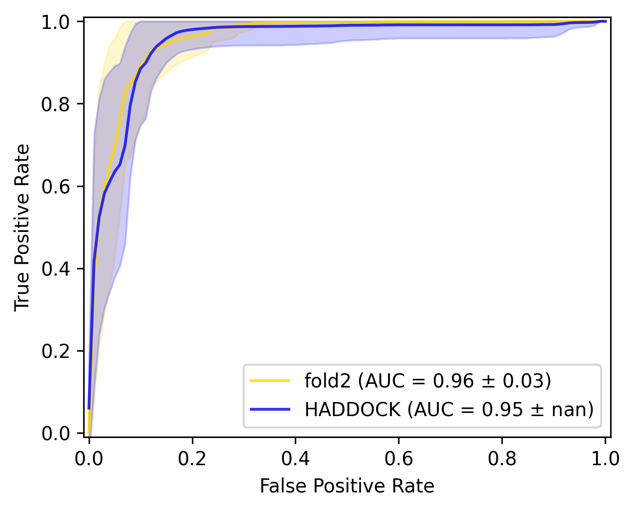

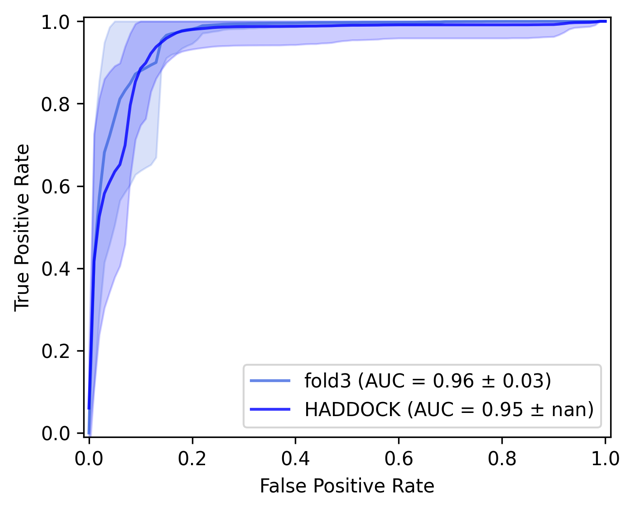

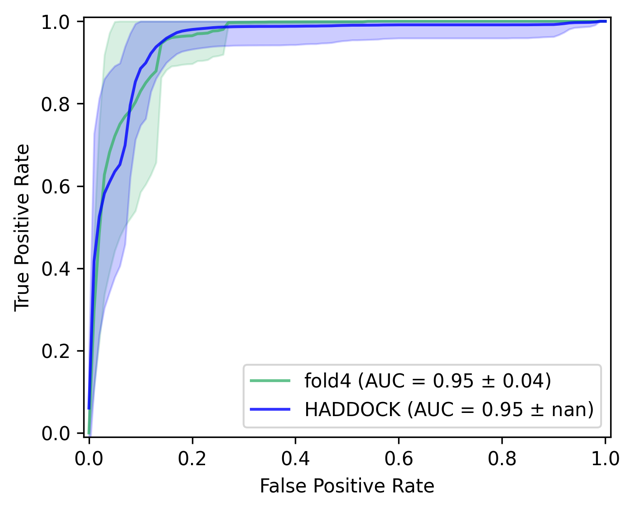

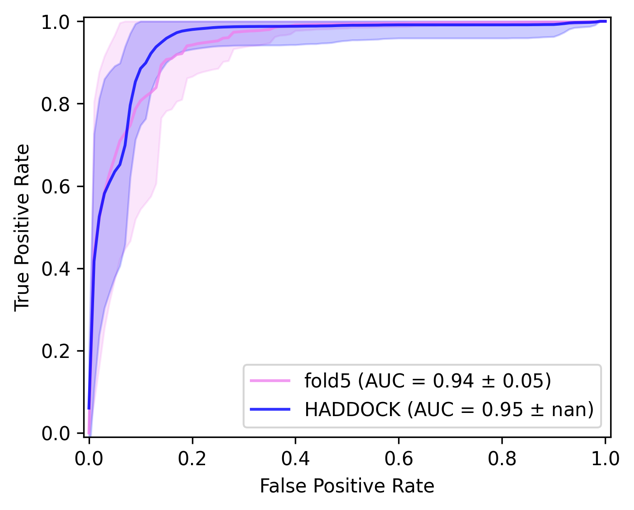

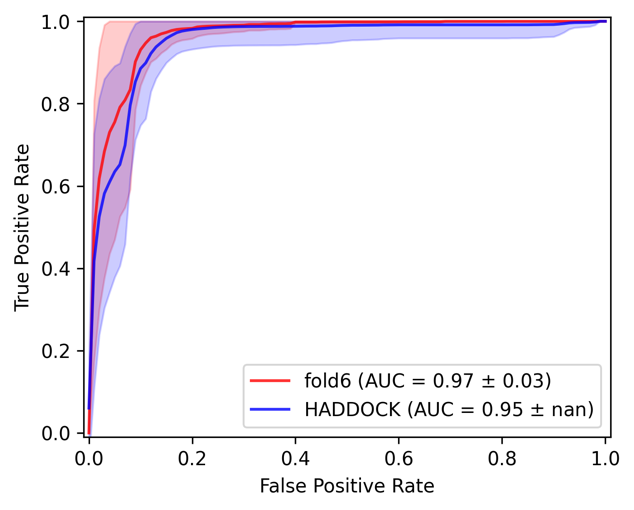

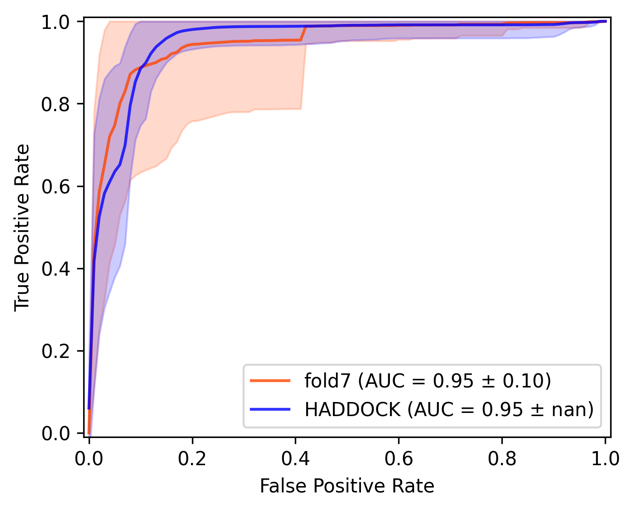

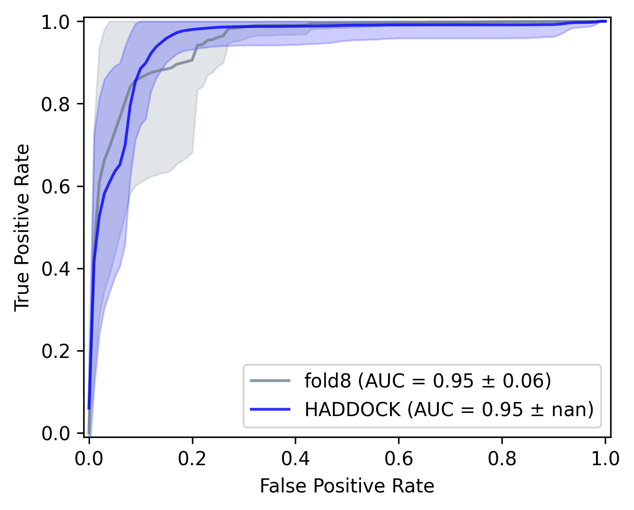

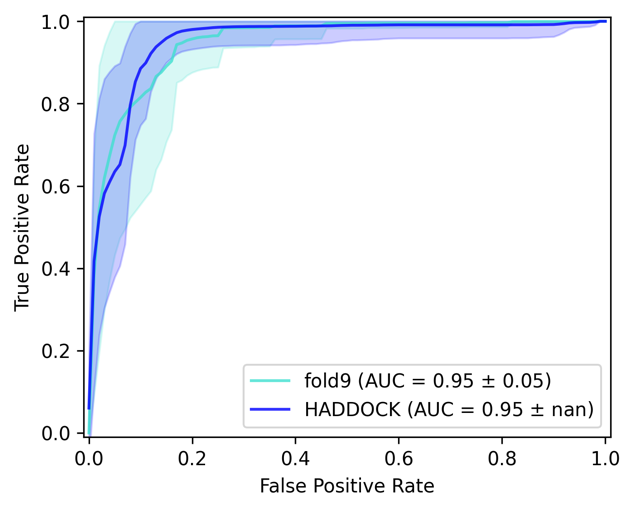

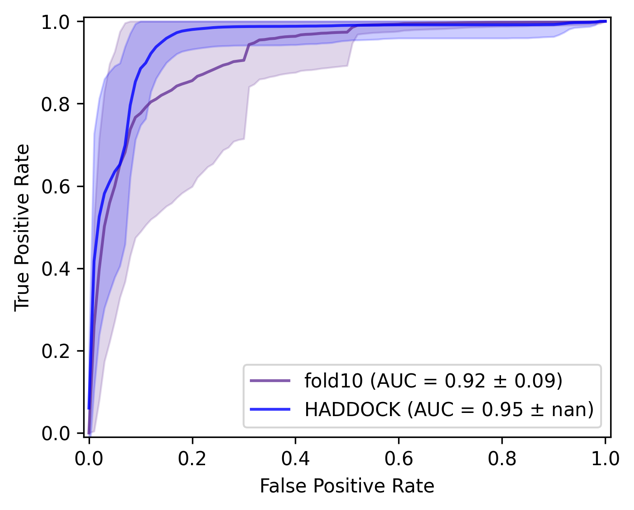

**Figure S5:** Hitrate obtained with the models retained for each DeepRank-GNN fold and HADDOCK score on each complex from the test set. A true positive case corresponds to a complex with fnat ≥ 0.3 correctly predicted.

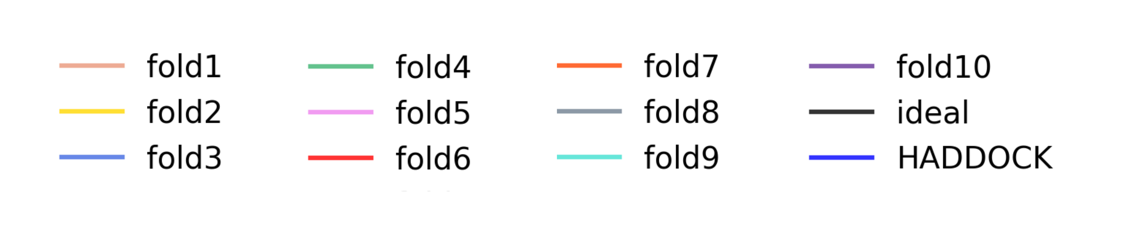

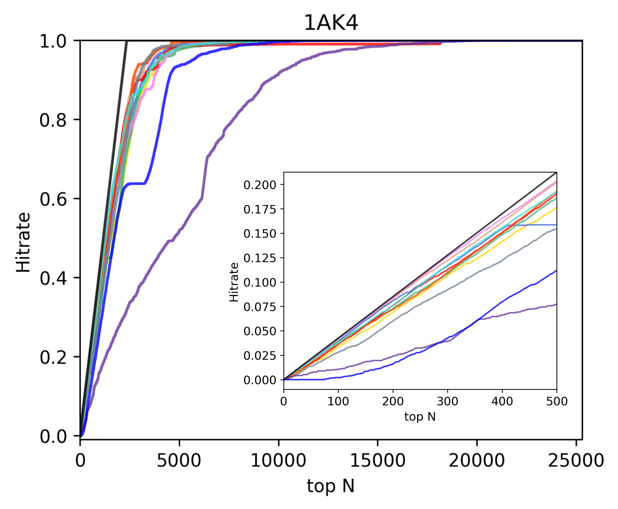

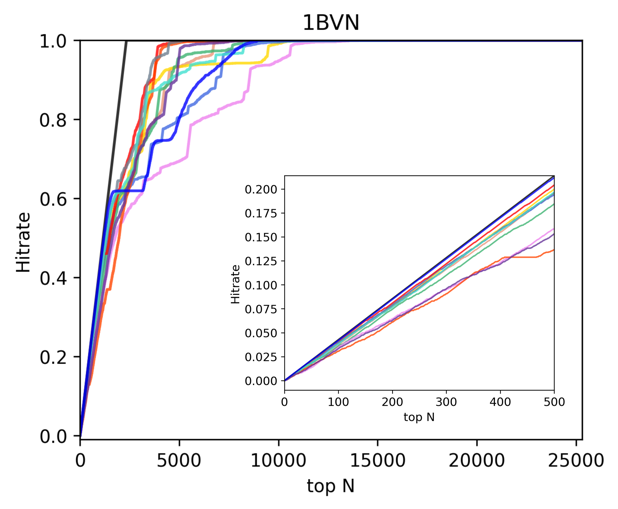

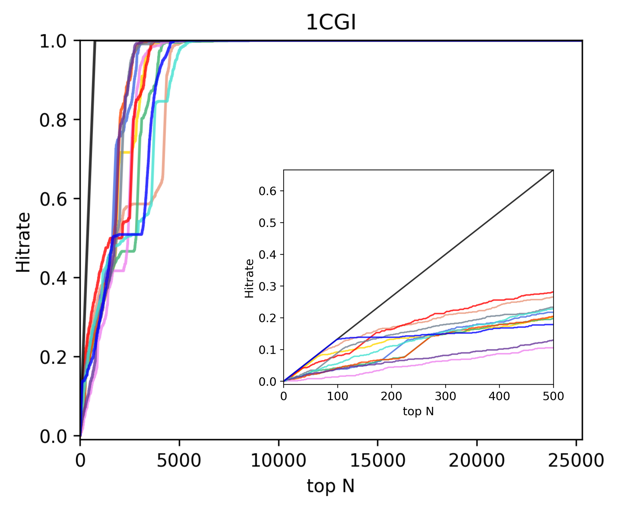

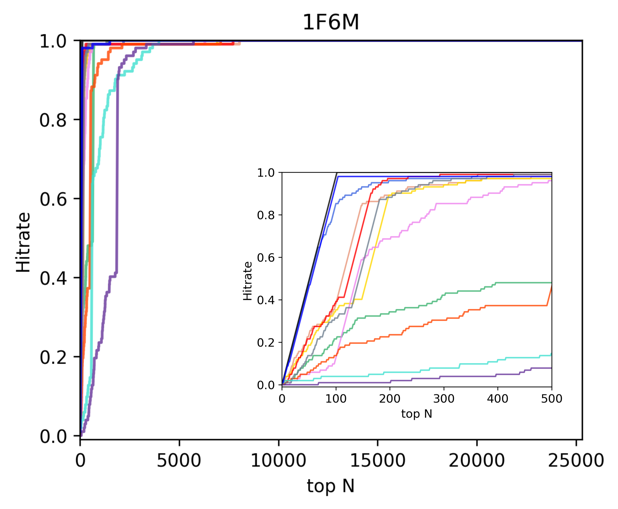

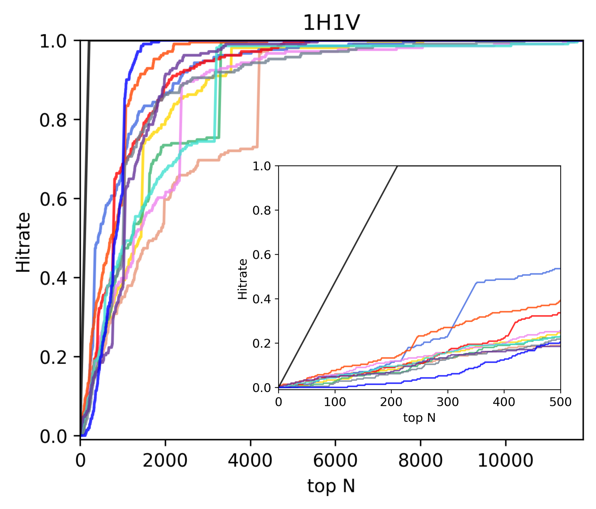

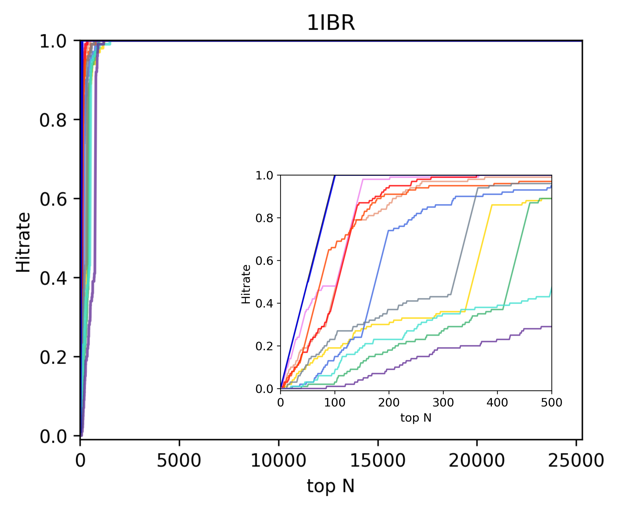

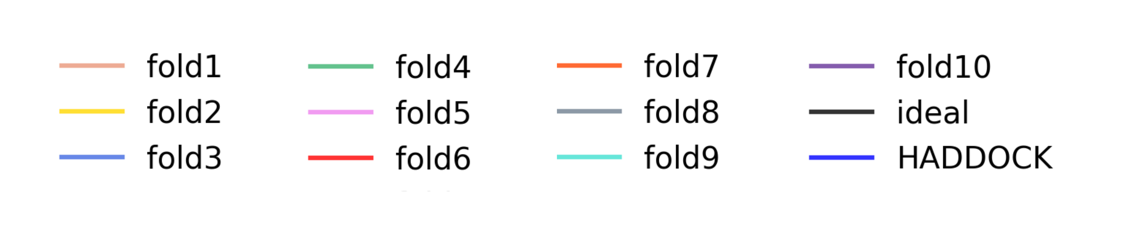

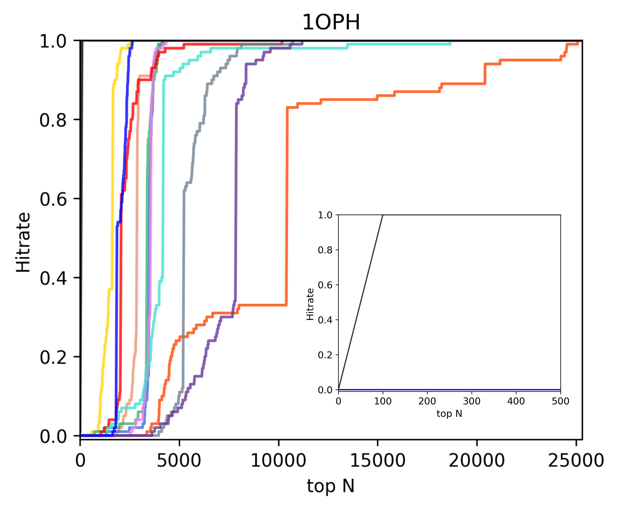

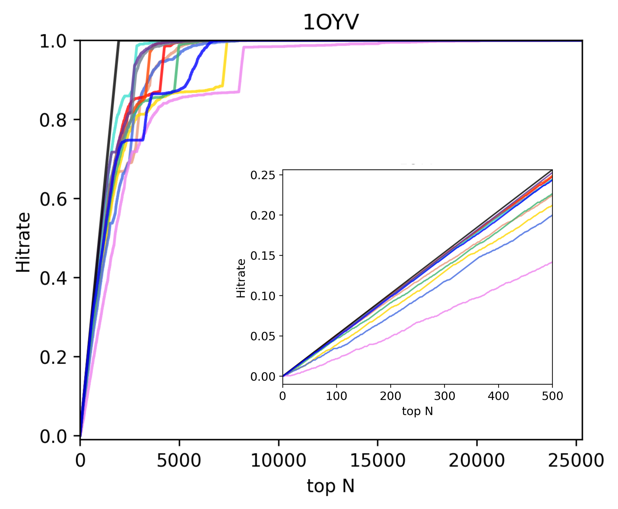

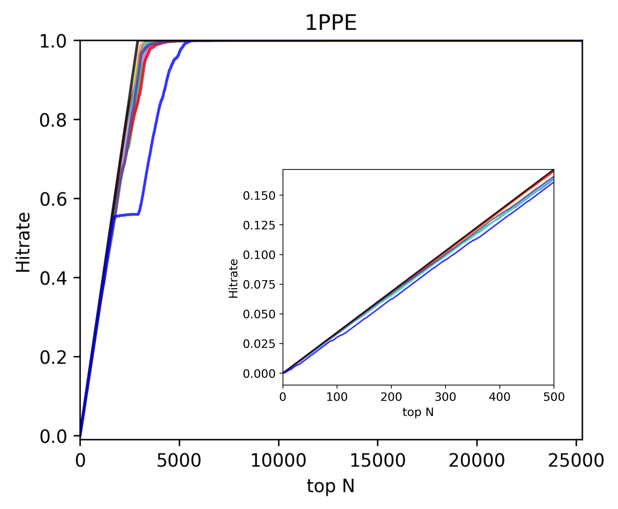

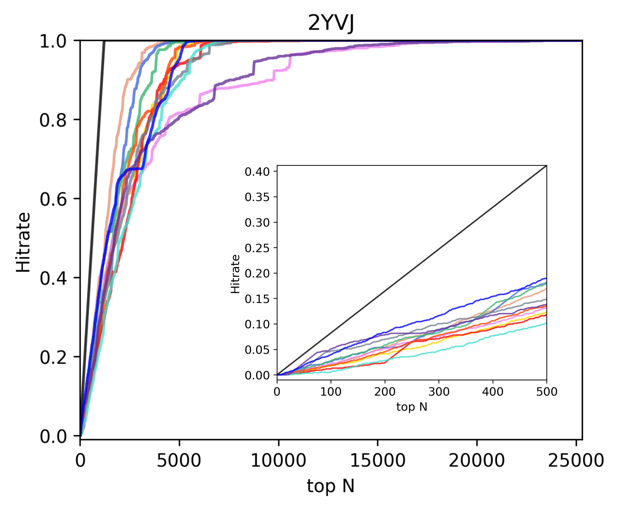

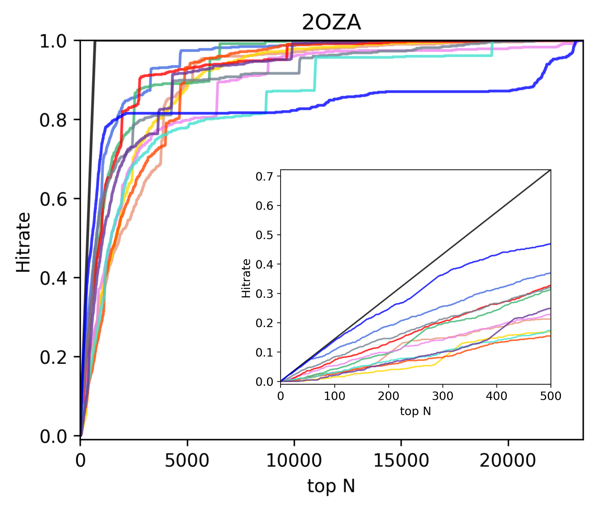

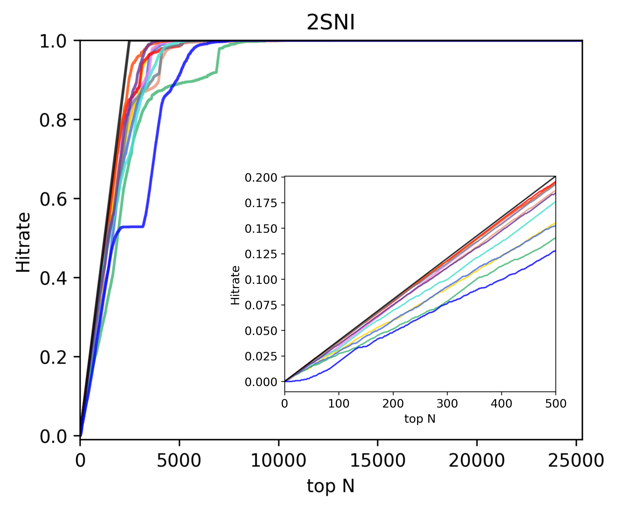

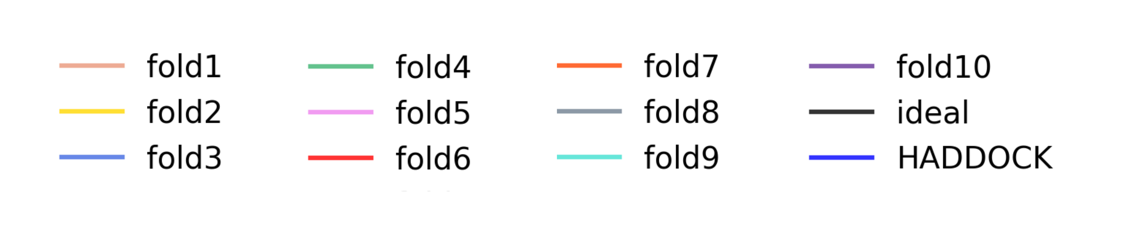

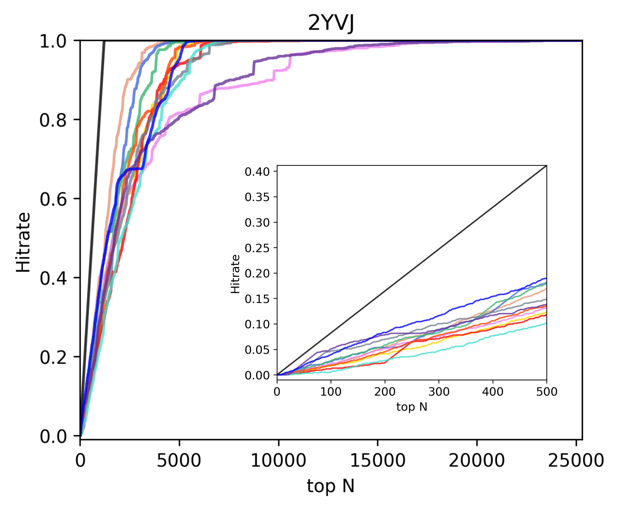

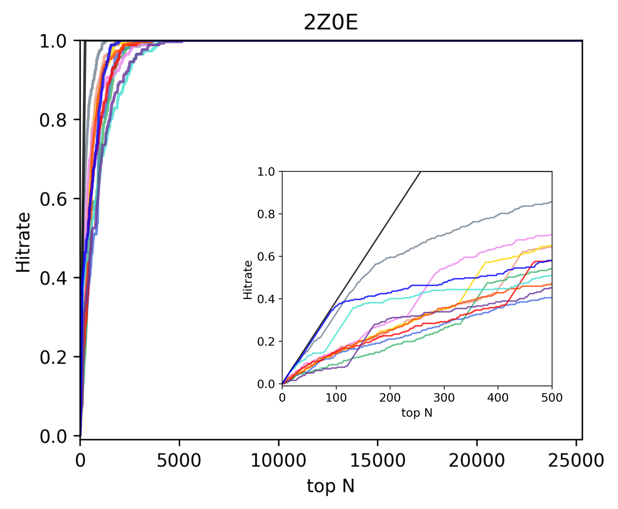

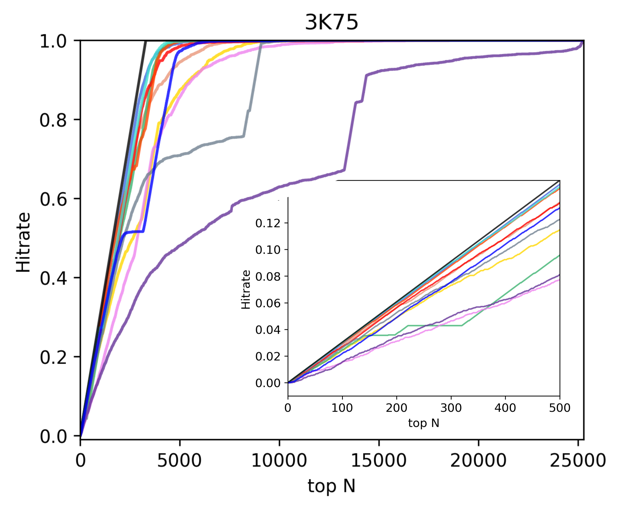

**Figure S6:** Correlation plots of the measured fnat (target) and the DeepRank-GNN score (prediction) (it0/it1/itw). The color code provides indications in the number of models associated to a plot area. The Spearman (rank) correlation is provide for each complex.

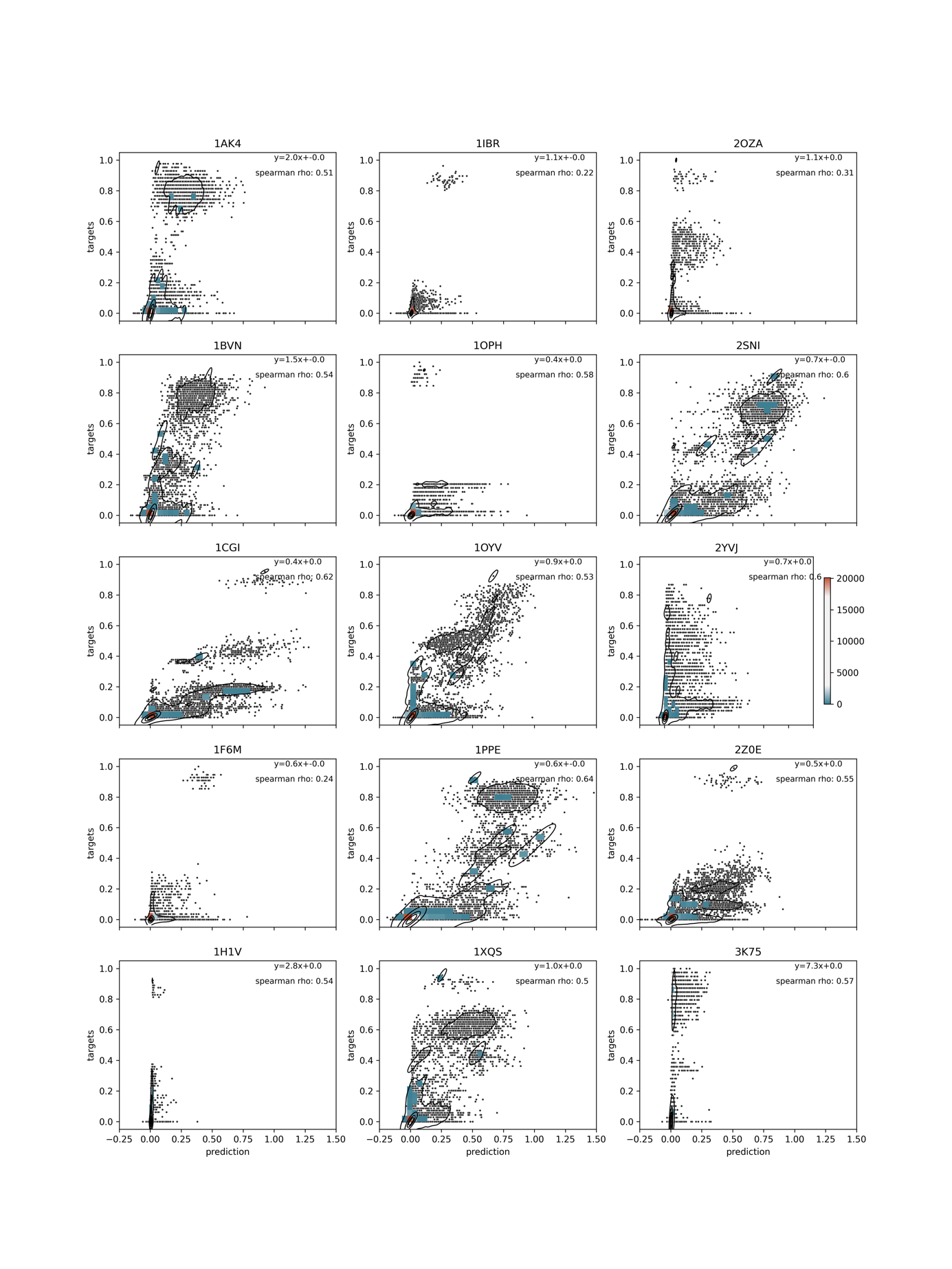

**Figure S7**: Correlation plots of the measured fnat (target) and the DeepRank-GNN score (prediction) (it1 and itw). The color code provides indications in the number of models associated to a plot area. The Spearman (rank) correlation is provide for each complex.

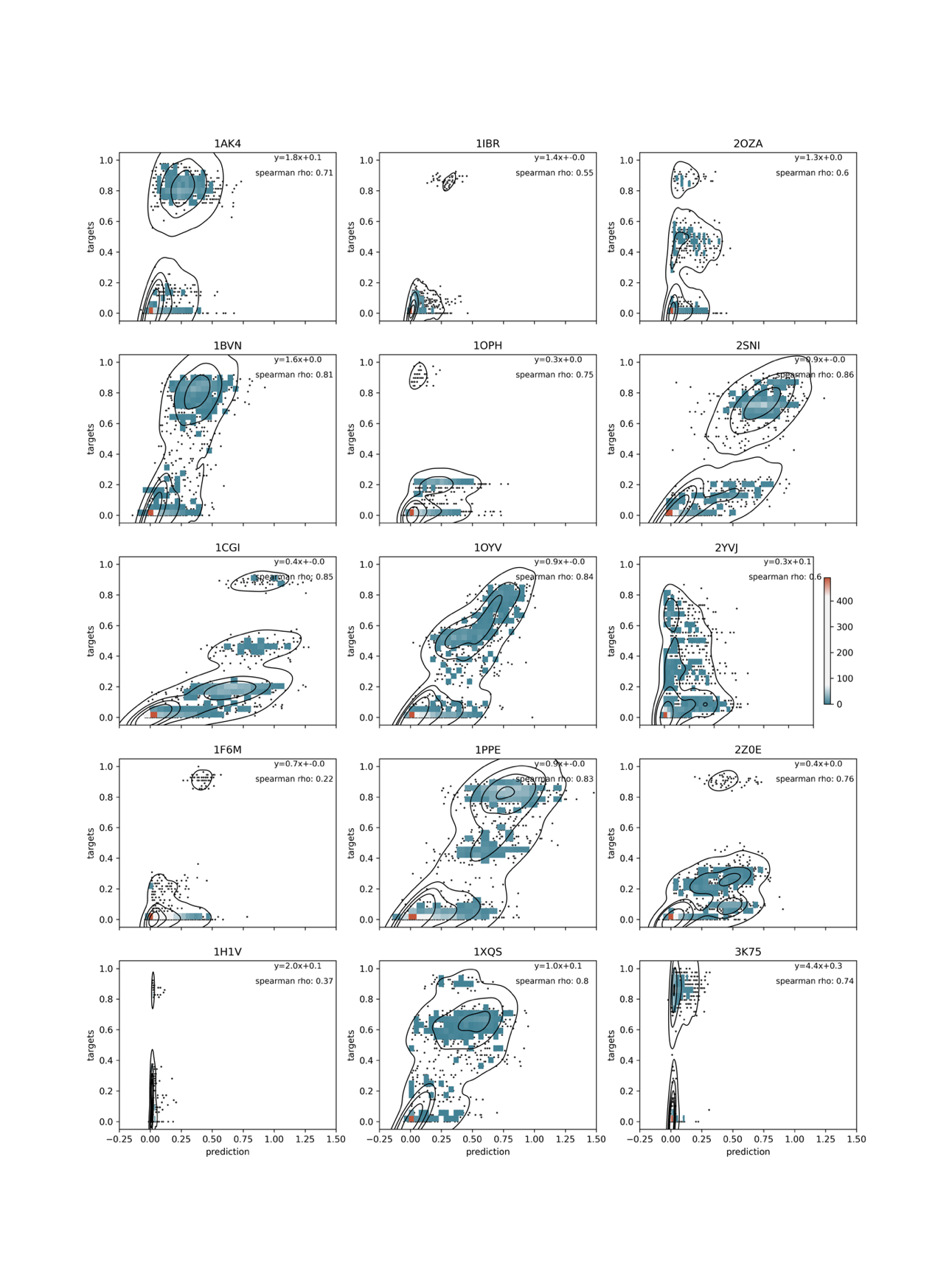

**Table S13**: Performance of the **graph** generation step of **DeepRank-GNN** on the 13 complexes (16666models) of the CAPRI score set using MPI distributed processes (4 CPUs).

|  |  |  |  |  |  |  |  | memory | **storage** | | | | |
| --- | --- | --- | --- | --- | --- | --- | --- | --- | --- | --- | --- | --- | --- |
|  | **number of model** | **time (s)** | **time/model (s)** | **diff=(moyenne-s)** | **diff*diff** | **(diff**2) * number model** | **time (min)** | **Maximum resident set size of the process during its lifetime, in Kbytes** | **Total in MB** | **MB/model** | **diff=(moyenne-MB)** | **diff*diff** | **(diff**2) * number model** |
| **T29** | 1979 | 1895,16 | 0,96 | -0,31 | 0,0933 | 184,71 | 31,59 | 1080508 | 297 | 0,15 | -0,01 | 0,0001 | 0,28 |
| **T30** | 1148 | 377,28 | 0,33 | 0,32 | 0,1046 | 120,13 | 6,29 | 1079676 | 118 | 0,10 | 0,04 | 0,0012 | 1,43 |
| **T32** | 599 | 499,83 | 0,83 | -0,18 | 0,0332 | 19,91 | 8,33 | 1077288 | 105 | 0,18 | -0,04 | 0,0014 | 0,83 |
| **T35** | 497 | 454,93 | 0,92 | -0,26 | 0,0693 | 34,44 | 7,58 | 1074596 | 77 | 0,15 | -0,02 | 0,0003 | 0,14 |
| **T37** | 1364 | 615,53 | 0,45 | 0,20 | 0,0403 | 55,03 | 10,26 | 1076640 | 184 | 0,13 | 0,00 | 0,0000 | 0,01 |
| **T39** | 1295 | 938,80 | 0,72 | -0,07 | 0,0053 | 6,87 | 15,65 | 1076396 | 179 | 0,14 | -0,00 | 0,0000 | 0,00 |
| **T40** | 1987 | 2381,25 | 1,20 | -0,55 | 0,2984 | 592,97 | 39,69 | 1076640 | 288 | 0,14 | -0,01 | 0,0000 | 0,09 |
| **T41** | 1101 | 321,28 | 0,29 | 0,36 | 0,1298 | 142,95 | 5,35 | 1074952 | 136 | 0,12 | 0,01 | 0,0002 | 0,23 |
| **T46** | 1570 | 700,97 | 0,45 | 0,21 | 0,0423 | 66,40 | 11,68 | 1077412 | 213 | 0,14 | 0,00 | 0,0000 | 0,01 |
| **T47** | 1015 | 319,02 | 0,31 | 0,34 | 0,1141 | 115,84 | 5,32 | 1085612 | 132 | 0,13 | 0,01 | 0,0001 | 0,07 |
| **T50** | 1447 | 1259,86 | 0,87 | -0,22 | 0,0478 | 69,11 | 21,00 | 1081640 | 205 | 0,14 | -0,00 | 0,0000 | 0,02 |
| **T53** | 1360 | 680,95 | 0,50 | 0,15 | 0,0229 | 31,19 | 11,35 | 1081852 | 204 | 0,15 | -0,01 | 0,0001 | 0,19 |
| **T54** | 1304 | 423,54 | 0,32 | 0,33 | 0,1071 | 139,72 | 7,06 | 1074460 | 164 | 0,13 | 0,01 | 0,0002 | 0,20 |
|  | **Total** | **Average time per model (s)** |  |  |  | **standard deviation s (per model)** |  |  | **average MB (per model)** |  |  |  | **standard deviation MB (per model)** |
|  | **16666** | **0,65** |  |  |  | **0,31** |  |  | **0,14** |  |  |  | **0,01** |

**Table S14**: Performance of the **grids** generation step of **DeepRank** on the 13 complexes (16666models) of the CAPRI score set using MPI distributed processes (4 CPUs) with no rotation of the input model.

|  |  |  |  |  |  |  |  | **memory** | **storage** | | | | |
| --- | --- | --- | --- | --- | --- | --- | --- | --- | --- | --- | --- | --- | --- |
|  | **number of model** | **time (s)** | **time/model (s)** | **diff=(moyenne-s)** | **diff*diff** | **(diff**2) * number model** | **time (min)** | **Maximum resident set size of the process during its lifetime, in Kbytes** | **Total in MB** | **MB/model** | **diff=(moyenne-MB)** | **diff*diff** | **(diff**2) * number model** |
| **T29** | 1979 | 31495,00 | 15,91 | -3,51 | 12,3547 | 24449,98 | 524,92 | 267880 | 7168 | 3,62 | -0,55 | 0,3024 | 598,45 |
| **T30** | 1148 | 10507,00 | 9,15 | 3,25 | 10,5445 | 12105,14 | 175,12 | 200200 | 3072 | 2,68 | 0,40 | 0,1569 | 180,17 |
| **T32** | 599 | 9923,00 | 16,57 | -4,17 | 17,3578 | 10397,31 | 165,38 | 226952 | 2048 | 3,42 | -0,35 | 0,1203 | 72,09 |
| **T35** | 497 | 9742,00 | 19,60 | -7,20 | 51,8678 | 25778,32 | 162,37 | 232060 | 1024 | 2,06 | 1,01 | 1,0237 | 508,76 |
| **T37** | 1364 | 15682,00 | 11,50 | 0,90 | 0,8147 | 1111,25 | 261,37 | 207764 | 4096 | 3,00 | 0,07 | 0,0048 | 6,53 |
| **T39** | 1295 | 20193,00 | 15,59 | -3,19 | 10,1976 | 13205,94 | 336,55 | 228060 | 4096 | 3,16 | -0,09 | 0,0082 | 10,68 |
| **T40** | 1987 | 24196,00 | 12,18 | 0,22 | 0,0495 | 98,39 | 403,27 | 242236 | 6144 | 3,09 | -0,02 | 0,0004 | 0,79 |
| **T41** | 1101 | 9213,00 | 8,37 | 4,03 | 16,2556 | 17897,46 | 153,55 | 197180 | 3072 | 2,79 | 0,28 | 0,0795 | 87,51 |
| **T46** | 1570 | 16685,00 | 10,63 | 1,77 | 3,1410 | 4931,37 | 278,08 | 214740 | 5120 | 3,26 | -0,19 | 0,0357 | 56,10 |
| **T47** | 1015 | 9194,00 | 9,06 | 3,34 | 11,1659 | 11333,43 | 153,23 | 199948 | 3072 | 3,03 | 0,05 | 0,0021 | 2,10 |
| **T50** | 1447 | 24763,00 | 17,11 | -4,71 | 22,2186 | 32150,33 | 412,72 | 237288 | 5120 | 3,54 | -0,47 | 0,2174 | 314,54 |
| **T53** | 1360 | 14393,00 | 10,58 | 1,82 | 3,3000 | 4487,99 | 239,88 | 206192 | 4096 | 3,01 | 0,06 | 0,0036 | 4,95 |
| **T54** | 1304 | 10667,00 | 8,18 | 4,22 | 17,8039 | 23216,22 | 177,78 | 204936 | 3072 | 2,36 | 0,72 | 0,5131 | 669,05 |
|  | **Total** | **Average time per model (s)** |  |  |  | **standard deviation s (per model)** |  |  | **average MB (per model)** |  |  |  | **standard deviation MB (per model)** |
|  | **16666** | **12,40** |  |  |  | **3,30** |  |  | **3,07** |  |  |  | **0,39** |

**Table S15**: Performance of the **grids** generation step of **DeepRank** on the 13 complexes (16666models) of the CAPRI score set using MPI distributed processes (4 CPUs) with 5 rotation of the input model, i.e. 6 orientation per model in total.

|  |  |  |  |  |  |  |  | **memory** | **storage** | | | | |
| --- | --- | --- | --- | --- | --- | --- | --- | --- | --- | --- | --- | --- | --- |
|  | **number of model** | **time (s)** | **time/model (s)** | **diff=(moyenne-s)** | **diff*diff** | **(diff**2) * number model** | **time (min)** | **Maximum resident set size of the process during its lifetime, in Kbytes** | **Total in MB** | **MB/model** | **diff=(moyenne-MB)** | **diff*diff** | **(diff**2) * number model** |
| **T29** | 1979 | 50674,00 | 25,61 | -1,66 | 2,7709 | 5483,62 | 844,57 | 283284 | 43008 | 21,73 | -2,19 | 4,8114 | 9521,73 |
| **T30** | 1148 | 26084 | 22,72 | 1,22 | 1,4884 | 1708,69 | 434,73 | 210112 | 18432 | 16,06 | 3,48 | 12,1310 | 13926,34 |
| **T32** | 599 | 23011 | 38,42 | -14,47 | 209,5093 | 125496,05 | 383,52 | 239180 | 13312 | 22,22 | -2,69 | 7,2092 | 4318,34 |
| **T35** | 497 | 21927 | 44,12 | -20,18 | 407,1297 | 202343,45 | 365,45 | 227408 | 11264 | 22,66 | -3,13 | 9,7674 | 4854,39 |
| **T37** | 1364 | 31326,00 | 22,97 | 0,97 | 0,9506 | 1296,60 | 522,10 | 223444 | 24576 | 18,02 | 1,52 | 2,3138 | 3155,97 |
| **T39** | 1295 | 34766,00 | 26,85 | -2,91 | 8,4395 | 10929,10 | 579,43 | 236760 | 26624 | 20,56 | -1,02 | 1,0412 | 1348,30 |
| **T40** | 1987 | 40965,00 | 20,62 | 3,32 | 11,0540 | 21964,23 | 682,75 | 249468 | 39936 | 20,10 | -0,56 | 0,3135 | 622,99 |
| **T41** | 1101 | 24444 | 22,20 | 1,74 | 3,0263 | 3331,94 | 407,40 | 213892 | 19456 | 17,67 | 1,87 | 3,4875 | 3839,77 |
| **T46** | 1570 | 32386,00 | 20,63 | 3,31 | 10,9775 | 17234,69 | 539,77 | 206932 | 30720 | 19,57 | -0,03 | 0,0008 | 1,25 |
| **T47** | 1015 | 24895 | 24,53 | -0,59 | 0,3432 | 348,35 | 414,92 | 201952 | 19456 | 19,17 | 0,37 | 0,1371 | 139,13 |
| **T50** | 1447 | 39310,00 | 27,17 | -3,23 | 10,4025 | 15052,45 | 655,17 | 246148 | 29696 | 20,52 | -0,98 | 0,9678 | 1400,38 |
| **T53** | 1360 | 29363,00 | 21,59 | 2,35 | 5,5263 | 7515,82 | 489,38 | 213712 | 25600 | 18,82 | 0,72 | 0,5115 | 695,60 |
| **T54** | 1304 | 19854,00 | 15,23 | 8,72 | 75,9651 | 99058,53 | 330,90 | 211440 | 23552 | 18,06 | 1,48 | 2,1826 | 2846,07 |
|  | **Total** | **Average time per model (s)** |  |  |  | **standard deviation s (per model)** |  |  | **average MB (per model)** |  |  |  | **standard deviation MB (per model)** |
|  | **16666** | **23,94** |  |  |  | **5,54** |  |  | **19,54** |  |  |  | **1,67** |

**Table S16:** Comparison of the computational performance of DeepRank-GNN and DeepRank in the training/evaluation phase using MPI distributed processes (4 CPUs). 80% of the 16666 CAPRI models fall into the training set, 20% into the evaluation set.

|  | **Number of epochs** | **data augmentation** | **Total elapsed time (in seconds)** | **Total elapsed time (in min)** | **Average elapsed time per epoch (in min)** | **Maximum resident set size of the process during its lifetime, in Kbytes** | **Maximum resident set size of the process during its lifetime, in Gbytes** |
| --- | --- | --- | --- | --- | --- | --- | --- |
| **deeprank-gnn** | 10,00 | None | 3458,23 | 57,64 | 5,76 | 1222200,00 | 1,17 |
| **deeprank (cnn)** grid size= (30,30,30) |  | None | 85752,41 | 1429,21 | 142,92 | 2106532,00 | 2,01 |
|  |  | 5 | 379651,62 | 6327,53 | 632,75 | 2211720,00 | 2,11 |

**Table S17:** DeepRank and DeepRank-GNN default features. The residue-level features highlighted in bold characters have been considered to train DeepRank and DeepRank-GNN in the comparative study detailed in section 3.3

|  | Name of the features | Number of parameters |
| --- | --- | --- |
| **DeepRank** | AtomicFeature | 6 |
|  | **FullPSSM** | **40** |
|  | **PSSM_IC** | **2** |
|  | **BSA** | **2** |
|  | **ResidueDensity** | **14** |
|  | atomicdensities | 8 |
|  | Total | 72 |
| **DeepRank-GNN** | **type** | **20** |
|  | **charge** | **1** |
|  | **polarity** | **4** |
|  | **BSA** | **1** |
|  | **PSSM** | **20** |
|  | **cons** | **1** |
|  | **ic** | **1** |
|  | **Total** | **48** |
